# BREC: An R package/Shiny app for automatically identifying heterochromatin boundaries and estimating local recombination rates along chromosomes

**DOI:** 10.1101/2020.06.29.178095

**Authors:** Yasmine Mansour, Annie Chateau, Anna-Sophie Fiston-Lavier

## Abstract

**Motivation:** Meiotic recombination is a vital biological process playing an essential role in genomes structural and functional dynamics. Genomes exhibit highly various recombination profiles along chromosomes associated with several chromatin states. However, eu-heterochromatin boundaries are not available nor easily provided for non-model organisms, especially for newly sequenced ones. Hence, we miss accurate local recombination rates, necessary to address evolutionary questions.

**Results:** Here, we propose an automated computational tool, based on the Marey maps method, allowing to identify heterochromatin boundaries along chromosomes and estimating local recombination rates. Our method, called **BREC** (heterochromatin **B**oundaries and **REC**ombination rate estimates) is non-genome-specific, running even on non-model genomes as long as genetic and physical maps are available. BREC is based on pure statistics and is data-driven, implying that good input data quality remains a strong requirement. Therefore, a data pre-processing module (data quality control and cleaning) is provided. Experiments show that BREC handles different markers density and distribution issues. BREC’s heterochromatin boundaries have been validated with cytological equivalents experimentally generated on the fruit fly *Drosophila melanogaster* genome, for which BREC returns congruent corresponding values. Also, BREC’s recombination rates have been compared with previously reported estimates. Based on the promising results, we believe our tool has the potential to help bring data science into the service of genome biology and evolution. We introduce BREC within an R-package and a Shiny web-based user-friendly application yielding a fast, easy-to-use, and broadly accessible resource.

**Availability:** BREC R-package is available at the GitHub repository https://github.com/ymansour21/BREC.

## Introduction

Meiotic recombination is a vital biological process which plays an essential role for investigating genome-wide structural as well as functional dynamics. Recombination events are observed in almost all eukaryotic genomes. Crossover, a one-point recombination event, is the exchange of DNA fragments between sister chromatids during meiosis. Recombination is a fundamental process that ensures genotypic and phenotypic diversity. Thereby, it is strongly related to various genomic features such as gene density, repetitive DNA, and DNA methylation [1–3].

Recombination rate varies not only between species, but also within species and within chromosomes. Different heterochromatin regions exhibit different profiles of recombination events. Therefore, in order to understand how and why recombination rate varies, it is important to break down the chromosome structure to smaller blocks where several genomic feature besides, recombination rate, are known to also exhibit different profiles. Chromatin boundaries allow to distinguish between two main states of chromatin that can be defined as euchromatin, which is lightly compact with a high gene density, and on the contrary, heterochromatin, which is highly compact with a paucity in genes. The heterochromatin is represented in different chromosome regions: the centromere and the telomeres. Euchromatin and heterochromatin regions exhibit different behaviours in terms of genomic features and dynamics related to their biologic function such as the cell division process that insures the organism viability. Consequently, easily distinguishing chromatin states is necessary for conducting further studies in various research fields and to be able to address questions related to cell processes such as: meiosis, gene expression, epigenetics, DNA methylation, natural selection and evolution, genome architecture and organization among others [4–6]. In particular, a profound understanding of centromeres, their complete and precise structure, organization and evolution is currently a hot research area. These repeat-rich heterochromatin regions are currently still either poorly or not assembled at all across eukaryote genomes. Despite the huge advances offered by NGS technologies, centromeres are still considered as enigmas, mostly because they are preventing genome assembly algorithms form reaching their optimal performance in order to achieve more complete whole genome sequences [7]. In addition, the highly diverse mechanisms of heterochromatin positioning [8] and repositioning [9] remain a complicated obstacle in face of fully understanding genome organization. Thus, generating high resolution genetic, physical and recombination maps, and locating heterochromatin regions is increasingly interesting the community across a large range of taxa [10–16].

Numerous methods for estimating recombination rates exist. Population genetic based-methods [17] provide accurate fine-scale estimates. Nevertheless, these methods are very expensive, time-consuming, require a strong expertise and, most of all, are not applicable on all kinds of organisms. Moreover, the sperm-typing method [18], which is also extremely accurate, providing high-density recombination maps, is male-specific and is applicable only on limited genome regions. On the other hand, a purely statistical approach, the Marey Maps [19], could avoid some of the above issues based on other available genomic data: the genetic and physical distances of genomic markers.

The Marey maps approach consists in correlating the physical map with the genetic map representing respectively physical and genetic distances for a set of genetic markers on the same chromosome. Despite the efficiency of this approach and mostly the availability of physical and genetic maps, generating recombination maps rapidly and for any organism is still challenging. Hence, the increasing need of an automatic, portable and easy-to-use solution.

Some Marey map-based tools already exist, two of which are largely used: (1) the MareyMap Online [20, 21] which is applicable on multiple species, however, it does not allow accurate estimate of recombination rates on specific regions like the chromosome extremities, and (2) the *Drosophila melanogaster* Recombination Rate Calculator (RRC) [22] which solves the previous issue by adjusting recombination rate estimates on such chromosome regions, yet, as indicated by its name, the RRC is *D. melanogaster-specifìc*. With the emerging Next Generation Sequencing (NGS) technologies, accessing whole chromosome sequences has become more and more possible on a wide range of species. Therefore, we may expect an exponential increase in markers number which will require more adapted tools to better handle such new scopes of data.

Here, we propose a new Marey map-based method as an automated computational solution that aims to, firstly, identify heterochromatin boundaries (HCB) along chromosomes, secondly, estimate local recombination rates, and lastly, adjust recombination rates on chromosome along the chromosomal regions marked by the identified boundaries. Our proposed method, called **BREC** (heterochromatin **B**oundaries and **REC**ombination rate estimates), is provided with an R-package and a Shiny web-based graphical user interface. BREC takes as input the same genomic data, genetic and physical distances, as in previous tools. It follows a workflow that, first, tests the data quality and offers a cleaning option, then, estimates local recombination rates and identify HCB. Finally, BREC re-adjusts recombination rate estimates along heterochromatin regions, the centromere and telomere(s), in order to keep the estimates as authentic as possible to the biological process [23]. Identifying the boundaries delimiting euchromatin and heterochromatin allows investigating recombination rate variations along the whole genome, which will help comparing recombination patterns within and between species. Furthermore, such functionality is fundamental for identifying the position of the centromeric and telomeric regions. Indeed, the position of the centromere on the chromosome has an influence on the chromatin environment and recent studies are interested in investigating how genome architecture may change with centromere organization [7].

Our results have been validated with cytological equivalents, experimentally generated on the fruit fly *D. melanogaster* genome [4, 24, 25]. Moreover, since BREC is non genome-specific, it could efficiently been run on other model as well as non-model organisms for which both genetic and physical maps are available. Even though it is still an ongoing study, BREC have also been tested with different further species and results are reported.

This paper is organized as follows: in the next Section (2), the new approach BREC is presented following a detailed step by step workflow. Section (3), presents the set of our results, based on both simulated and real data. The results are then discussed in Section (4). Concluding remarks with some perspectives are outlined in Section (5). Further details of the data involved, how the methods were calibrated and validated are reported within the supplementary materials.

## New Approach: BREC

BREC is designed following the workflow represented in Fig 1. In order to ensure that the widest range of species could be analyzed by our tool, we designed a pipeline which adapts behaviour with respect to input data. Mostly, each step of the pipeline relies on statistical analysis, adaptive algorithms and decision proposals led by empirical observation.

The workflow starts with a pre-processing module (called “Step 0”) aiming to prepare the data prior to the analysis. Then, it follows six main steps: (1) estimate Marey Map-based local recombination rates, (2) identify chromosome type, (3) prepare the HCB identification, (4) identify the centromeric boundaries, (5) identify the telomeric boundaries, and (6) extrapolate the local recombination rate map and generate an interactive plot encompassing all BREC outputs (see Fig 1). Each step is detailed hereafter.

**Fig 1.**
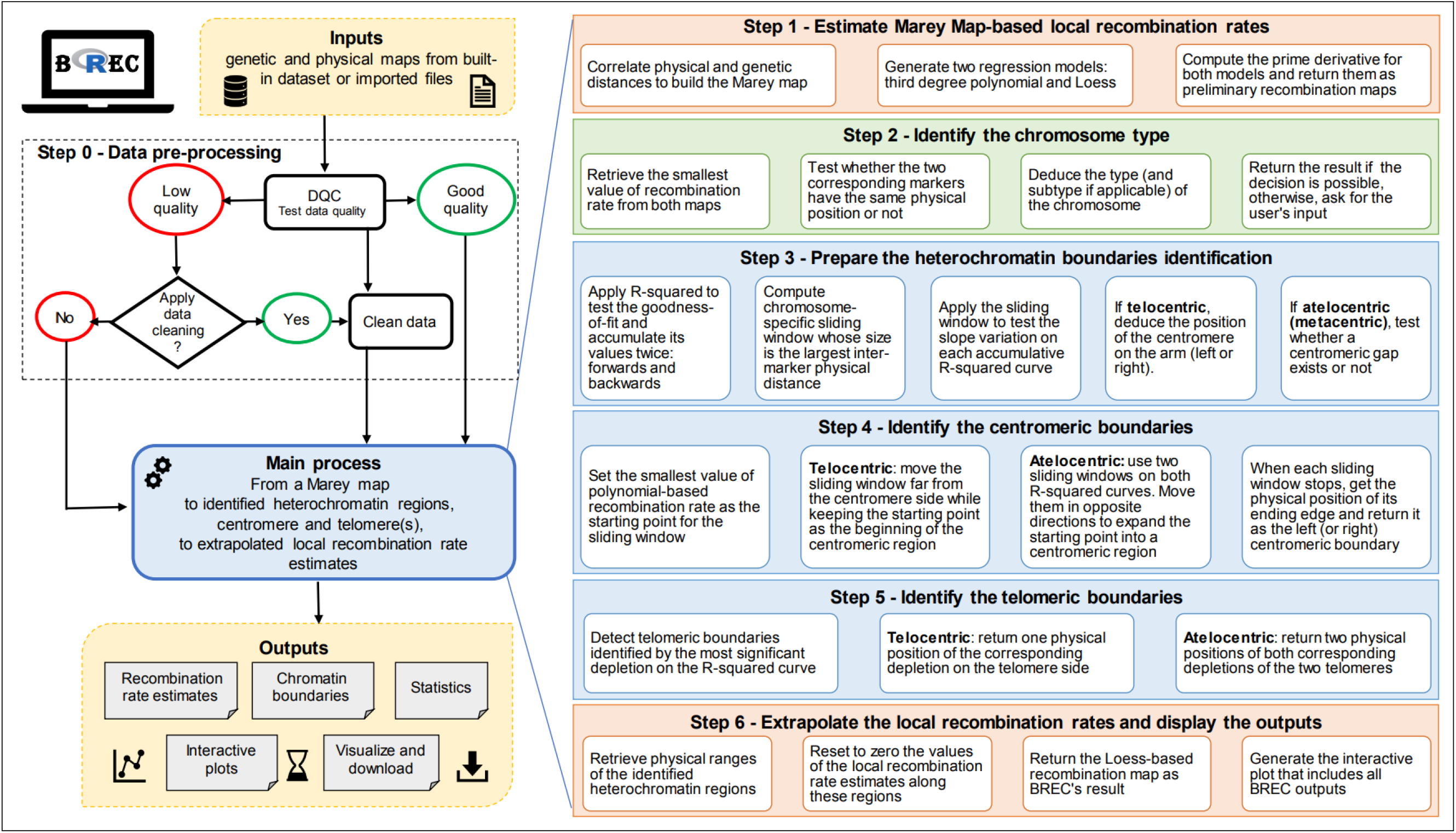
BREC workflow. This figure provides an overview of the tool design explaining how the different modules are linked together and how BREC functionalities are implemented. The diagram starting from the top left side shows input data, how are they processed, and what outputs are expected to be returned. Then, the right part of the figure, representing a zoom-in on BREC’s main module (estimating recombination rates, identifying chromosome type, identifying HCB, extrapolating the recombination map and generating the interactive plot), clarifies each step following a more detailed scheme.

### Step 0 - Apply data pre-processing

Since we have noticed that BREC estimates are sensitive to the quality of input data, we propose a pre-processing step to assess data quality and suggest an optional data cleaning for outliers. As such, we could ensure a proper functioning during further steps.

#### Data quality control

The quality of input data is tested regarding two criteria: (1) density of markers and (2) the homogeneity of their distribution on the physical map, along a given chromosome. First, the mean density, defined as the number of markers per physical map length, is computed. This value is compared with the minimum required threshold of 2 markers/Mb. Based on the displayed results, the user gets to decide if data cleaning is required or not. The threshold of 2 markers/Mb is selected based on a simulation process that allowed to test BREC results while decreasing markers density until the observed HCB estimates seemed to be no longer exploitable (see Materials and Methods in Section). Second, the distribution of input data is tested *via* a comparison with a simulated uniform distribution of identical markers density and physical map length. This comparison is applied using Pearson’s *Chi – squared* test [26] which allows to examine how close the observed distribution (input data) is to the expected one (simulated data).

#### Data cleaning

The cleaning step aims to reduce the disruptive impact of noisy data, such as outliers, in order to provide more accurate recombination rate and heterochromatin boundary results. If the input data fails to pass the Data Quality Control (DQC) test, the user has the option to apply or not a cleaning process. This process consists of identifying the extreme outliers and eliminating them upon the user’s confirmation. Outliers are detected using the distribution statistics of the genetic map (see Fig S1). More precisely, inter-marker distances (separating each two consecutive points) are computed along the genetic map. Using a boxplot, distribution statistics (quartiles, mean, median) are applied on these inter-marker distances in order to identify outliers, which are chosen as the 5% of the data points with a genetic distance greater than the maximum extreme value, and should be discarded. Thus, the cleaning is targeting markers for which the genetic distance is quite larger than most of the rest. After the first cleaning iteration, DQC is applied again to assess the new density and distribution. The user can also choose to bypass the cleaning step, but in such case, BREC’s behaviour is no longer guaranteed.

### Step 1 - Estimate Marey Map-based local recombination rates

Once the data are cleaned, the recombination rate can be estimated based on the Marey map [19] approach by: (1) correlating genetic and physical maps, (2) generating two regression models -third degree polynomial and Loess-that better fits these data, (3) computing the prime derivative for both models which will represent preliminary recombination maps for the chromosome. The main purpose of interpolation here is to provide local recombination rate estimates for any given physical position, instead of only the ones corresponding to available markers.

At this point, both recombination maps are used to identify the chromosome type as well as the approximate position of centromeric and telomeric regions. Yet, as a final output, BREC will return only the Loess-based adjusted map for recombination rates since it provides finer local estimates than the polynomial-based map.

### Step 2 - Identify chromosome type

BREC provides a function to identify the type of a given chromosome, with respect to the position of its centromere. This function is based on the physical position of the smallest value of recombination rate estimates, which primarily indicate where the centromeric region is more likely to be located. Our experimentation allowed to come up with the following scheme (see Fig S2). Two main types are identified: telocentric and atelocentric [27]. Atelocentric type could be either metacentric (centromere located approximately in the center with almost two equal arms) or not metacentric (centromere located between the center and one telomere of the chromosome). The latter includes the two most known subtypes, submetacentric and acrocentric (recently considered as types rather than subtypes). It is tricky for BREC to correctly distinguish between submetacentric and acrocentric chromosomes because the position of their centromeres varies slightly, and capturing this variation (based on the smallest value of recombination rate on both maps -polynomial and Loess-) could not be achieved, yet. Therefore, we chose to provide this result only if the identification process allowed to automatically identify the subtype. Otherwise, the user gets the statistics on the chromosome and is invited to decide according to further *a priori* knowledge. The two subtypes (metacentric and not metacentric) are distinguished following an intuitive reasoning inspired by their definition found in the literature. First, BREC identifies whether the chromosome is an arm (telocentric) or not (atelocentric). Then, test if the physical position of the smallest value of the estimated recombination rate is located between 40% and 60% interval, the subtype is displayed as *metacentric*, otherwise, it is displayed as not *metacentric*. The recombination rate is estimated using the Loess model (“LOcal regrESSion”) [28, 29].

### Step 3 - Prepare the HCB identification

The HCB identification is a purely statistical approach relying on the coefficient of determination *R*^2^, which measures how good the generated regression model fits the input data [30]. We chose this approach because the Marey map usually exhibits lower quality of markers (density and distribution) on the heterochromatin regions. Thus, we aim to capture this transition from high to low quality regions (or *vice versa*) as it reflects the transition from euchromatin to heterochromatin regions (or *vice versa*). The coefficient *R*^2^ is defined as the cumulative sum of squares of differences between the interpolation and observed data. *R*^2^ values are accumulated along the chromosome. In order to eliminate the biased effect of accumulation, *R*^2^ is computed twice: *R*^2^ – *forward* starts the accumulation from the beginning of the chromosome to provide the left centromeric and left telomeric boundaries, while *R*^2^ – *backwards* starts from the end of the chromosome providing the right centromeric and right telomeric boundaries. These *R*^2^ values were calculated using the `rsq` package in R. To compute *R*^2^ cumulative vectors, `rsq` function is applied on the polynomial regression model. In fact, there is no such function for non-linear regression like Loess, because in such models, high *R*^2^ does not always mean good fit. A sliding window is defined and applied on the *R*^2^ vectors with the aim of precisely analysing their variations (see details in the next step). In case of a telocentric chromosome, the position of the centromere is then deduced as the left or the right side of the arm, while in case of an atelocentric chromosome, the existence of a centromeric gap is investigated.

### Step 4 - Identify centromeric boundaries

Since the centromeric region is known to present reduced recombination rates, the starting point for detecting its boundaries is the physical position corresponding to the smallest polynomial-based recombination rate value. Then, a sliding window is applied in order to expand the starting point into a region based on *R^2^* variations in two opposite directions. The size of the sliding window is automatically computed for each chromosome as the largest value of ranges between each two consecutive positions on the physical map (indicated as *i* and *i* + 1 in Equation 1). After making sure the sliding window includes at least two data points, the mean of local growth rates inside the current window is computed and tested compared to zero. If it is positive (resp. negative) on the forward (resp. backwards) *R^2^* curve, the value corresponding to the window’s ending edge is returned as the left (resp. right) boundary. Else, the window moves by a step value equal to its size.

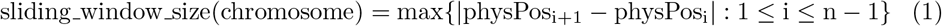

There are some cases where chromosome data present a centromeric gap. Such lack of data produces biased centromeric boundaries. To overcome this issue, chromosomes with a centromeric gap are handled with a slightly different approach: after comparing the mean of local rates of growth regarding to zero, accumulated slopes of all data points within the sliding window are computed adding one more point at a time. If the mean of accumulated slopes keeps the same variation direction as the mean of growth rates, the centromeric boundary is set as the ending edge of the window. Else, the window slides by the same step value as before (equal to its size). The difference between the two chromosome types is that for the telocentric case, only one sliding window is used, it’s starting point is the centromeric side, and it moves away from it. As for the atelocentric case, two sliding windows are used (one on each *R*^2^ curve), their starting point is the same, and they move in opposite directions to expand the centromere into a region.

### Step 5 - Identify telomeric boundaries

Since telomeres are considered heterochromatin regions as well, they also tend to exhibit a low fitness between the regression model and the data points. More specifically, the accumulated *R*^2^ curve tends to present a significant depletion around telomeres. Therefore, a telomeric boundary is defined here as the physical position of the most significant depletion corresponding to the smallest value of the *R*^2^ curve. As such, in the telocentric case, only one *R*^2^ curve is used and it gives one boundary of the telomeric region (the other boundary is defined by the beginning of the left telomere or the end of the right telomere). Whilst in the atelocentric case, where the are two telomeres, the depletion on *R*^2^ – *forward* detects the end of the left telomeric region and the depletion on *R*^2^ – *backwards* detects the beginning of the right telomeric region. The other two boundaries (the beginning of the left telomere and the end of the right telomere) are defined to be, respectively, the same values of the two markers with the smallest and the largest physical position available within the input data of the chromosome of interest.

### Step 6 - Extrapolate the local recombination rate estimates and generate interactive plot

The extrapolation of recombination rate estimates within the identified centromeric and telomeric regions automatically performs an adjustment by resetting the initial biased values to zero along these heterochromatin ranges. Then, each of the above BREC outputs are combined to generate one interactive plot displayed for visualisation and download (see details in Section).

## Results

In this section, we present the results obtained through the following validation process. First, we automatically re-identified HCB with approximate resolution to the reference equivalents. Second, we tested the robustness of BREC method according to input data quality, using the well-studied *D. melanogaster* genome data, for which recombination rate and HCB have already been accurately provided [4, 22, 24, 31](Fig S3). In addition, we extended the robustness test to a completely different genome, the domesticated tomato *S. lycopersicum* [32] to better interpret the study results. Even if the Loess span value does not impact the HCB identification, but only the resulting recombination rate estimates, the span values used in this study are: 15% for *D. melanogaster* (for comparison purpose) and 25% for the rest of experiments. Our analysis shows that BREC is applicable on data from a various range of organisms, as long as the data quality is good enough. BREC is data-driven, thus, the outputs are strongly dependant of the markers density, distribution and chromosome type specified (automatically, or with the user’s a priori knowledge).

### Approximate, yet congruent HCB

#### Fruit fly genome *D.melanogaster*

Our method for identifying HCB has been primarily validated with cytological data experimentally generated on the *D. melanogaster* Release 5 genome [4, 24, 25, 33]. For all five chromosomal arms (X, 2L, 2R, 3L, 3R). his genome presents a mean density of 5.39 markers/Mb and a mean physical map length of 22.92Mb. We obtained congruent HCB with a good overlap and shift, distance between the physical position of the reference and BREC, from 20Kb to 4.58Mb (see Section). We did not observe a difference in terms of mean shift for the telomeric and centromeric BREC identification (*χ*^2^ = 0.10, df = 1, *p − value* = 0.75)(See Tables 1 and S1). We observe a lower resolution for the chromosomal arms 3L and 3R (see Fig S4). This suggests that the data for those two chromosomal arms might not present a quality as good as the rest of the genome. Interestingly, the local markers density for these two chromosomal arms shows a high variation, not like for the other chromosomal arms. For instance, the 2L for which BREC returns accurate results, shows a lower variation (see Fig S5). Without these two arms, the max shift for both centromeric and telomeric BREC boundaries is smaller than 1.54Mb with a mean shift decreasing from 1.43Mb to 0.71Mb.

**Table 1.** BREC HCB compared to reference boundaries from the reference genome of *D. melanogaster*. The shift is the absolute value of the distance between the BREC and the reference physical heterochromatin boundary. The first five rows represent all chromosomal arms. Grouped columns present reference, BREC and shift values for the centromeric boundaries (Columns 2-4), and for the telomeric boundaries (Columns 4-6). Here the boundary values correspond to the internal HCB. The external boundaries are represented by the physical positions of the first and the last markers of the chromosomes. All values are expressed in Megabase (Mb). The red asterisk indicates the largest shift value reported on centromeric and telomeric boundaries separately (see corresponding Fig S4). The last four rows represent general statistics on the shift value. From top to bottom, they are minimum, maximum, mean, and median respectively. See details on the shift metrics in Section.

| Chromosomal arm | Centromeric (Mb) |  |  | Telomeric (Mb) |  |  |
| --- | --- | --- | --- | --- | --- | --- |
|  | Boundaries |  | Shift | Boundaries |  | Shift |
|  | Reference | BREC |  | Reference | BREC |  |
| X | 20.67 | 20.10 | 0.56 | 2.46 | 0.92 | 1.54 |
| 2L | 19.95 | 20.33 | 0.38 | 0.70 | 0.68 | 0.02 |
| 2R | 6.09 | 5.01 | 1.08 | 20.02 | 20.71 | 0.69 |
| 3L | 18.41 | 20.30 | 1.90 | 0.36 | 2.26 | 1.91* |
| 3R | 8.35 | 3.77 | 4.58* | 27.25 | 25.64 | 1.61 |
| Min. shift | 0.38 |  |  | 0.02 |  |  |
| Max. shift | 4.58 |  |  | 1.91 |  |  |
| Mean shift | 1.70 |  |  | 1.15 |  |  |
| Median shift | 1.08 |  |  | 1.54 |  |  |

This first analysis suggests that BREC method returns accurate results on this genome. However, the boundaries identification process appears very sensitive to the local density and distribution of the markers along a chromosome (see Fig S4). Therefore, we conducted further experiments on a different dataset, the tomato genome (see Fig S6).

#### Tomato genome *S. lycopersicum*

Results of experimenting BREC behaviour on all 12 chromosomes of *S. lycopersicum* genome [32] are shown as values in Table S3 and as plots in Fig S7. This genome presents a mean density of 2.64 markers/Mb and a mean physical map length of 62.71Mb. We observe a variation in the shift value representing the difference on the physical map between reference HCB and their equivalents returned by BREC. Unlike *D. melanogaster* genome which is of a smaller size, with five telocentric chromosomes (chromosomal arms) and a strongly different markers distribution, the tomato genome exhibits a completely different study case. This is a plant genome, with an approximately 8-fold bigger size genome. It is organized as twelve atelocentric chromosomes of a mean size of 60Mb except for chromosomes 2 and 6 which are more likely to be rather considered telocentric based on their markers distribution. Also, we observe a long plateau of markers along the centromeric region with a lower density than the rest of the chromosomes, something which highly differs from *D. melanogaster* data. We believe all these differences between both genomes gives a good validation but also evaluation for BREC behaviour towards various data quality scenarios. Furthermore, since BREC is a data-driven tool, these experiments help analysing data-related limitations that BREC could be facing while resolving differently. From another view point, BREC results on the tomato genome highlights the fact that markers distribution along heterochromatin regions, in particular, strongly impacts the identification of eu-heterochromatin boundaries, even when the density is of 2 markers/Mb or more.

### Consistency despite the low data quality

We aim in this part to study to what extent BREC results are depending on the data quality.

#### BREC handles low markers density

We start by assessing the marker density on the BREC estimates. We generated simulated datasets with decreasing fractions of markers for each chromosomal arms (from 100% to 30%). For that, we randomly select a fraction of markers 30 times and compute the mean shift between the BREC and the reference telomeric and centromeric boundaries. We note that BREC’s resolution decreases drastically with the fraction and thus with the marker density (see Fig S8). However, BREC results appears stable until 70% of the data for all the chromosomal arms and more specifically for the telomeric boundary detection. Only for the centromeric boundary of the chromosomal arm 3R, we observe the opposite pattern: BREC returns more accurate telomeric boundary estimates when the number of markers decreases. This supports the low quality of the data around the 3R centromere.

This simulation process allowed to set a min density threshold representing the minimum value for data density in order to guarantee an accurate results of BREC estimates at 5 markers/Mb (fraction of around 70% of the data) on average in *D. melanogaster*. This analysis also supports that as the marker density alone can not explain the BREC resolution, BREC may be also sensitive to the marker distribution.

Fig S5 clearly shows that markers density varies within and between the five chromosomal arms with a mean of 4 to 8 markers/Mb. The variance is induced by the extreme values of local density, such as 0 or 24 markers/Mb on the chromosomal arm X. Still, the overall density is around 5 markers/Mb for the whole genome.

#### BREC handles heterogeneous distribution

Along chromosomes, genetic markers are not homogeneously distributed. Therefore, to assess the impact of the markers distribution on BREC results, we designed different *data scenarios* with respect to reference data distribution (see Materials and Methods:). We choose as reference the chromosomal arms 2L and 2R of *D. melanogaster* as we obtained accurate results for these two chromosomal arms. After the concatenation of the two arms 2L and 2R, we ended up with a metacentric simulated chromosome as a starting simulation (total physical length of 44Mb). While this length was kept unchanged, markers local density and distribution were modified (see Materials and Methods:; Fig S9).

One particular yet common case is the centromeric gap. Throughout our analysis, we consider that a chromosome presents a centromeric gap if its data exhibit a lack of genetic markers on a relatively large region on the physical map. As centromeric regions usually are less accessible to sequence due to its high compact state. Consequently, these regions are also hard to assemble and that is why a lot of genomes have chromosomes presenting a centromeric gap. It is important to know that a centromeric gap is not always exactly located on the middle of a chromosome. Instead, its physical location depends on the type of chromosome (see more details on Fig S2).

We also assess the veracity of BREC on datasets with variable distribution using simulated data with and without centromeric gap (see Fig S9).

For all six simulation datasets, BREC results overlap the reference boundaries. Thus BREC correctly handles the presence of a centromeric gap (see Fig S9: (a)(c)(e)). BREC stays robust to a non-uniform distribution of markers, under the condition that regions bordering the boundaries are greater than 2 markers/Mb (see Fig S10). In case of non-uniform distribution, BREC resolution is higher when the local density is stronger around heterochromatin regions (see Fig S9: (c)(d)(e)(f)). This suggests that low density on euchromatin regions far from the boundaries is not especially a problem either.

### Accurate local recombination rate estimates

After the identification of HCB, BREC provides optimized local estimates of recombination rate along the chromosome by taking into account the absence of recombination in heterochromatin regions. Recombination rates are set to zero across the centromeric and telomeric regions regardless of the regression model. To closely compare the third degree polynomial with Loess, using different span values, we experimented this aspect on *D. melanogaster* chromosomal arms and reported the results in Fig S11.

To assess the veracity of the recombination rates along the whole genome, we compared BREC results with previous recombination rate estimates (see Fig 2; [4, 24]). BREC recombination rate estimates are significantly strongly correlated with reference data (Spearman’s: *P* << 0.001) while the reference estimates fail in telomeric regions.

**Fig 2.**
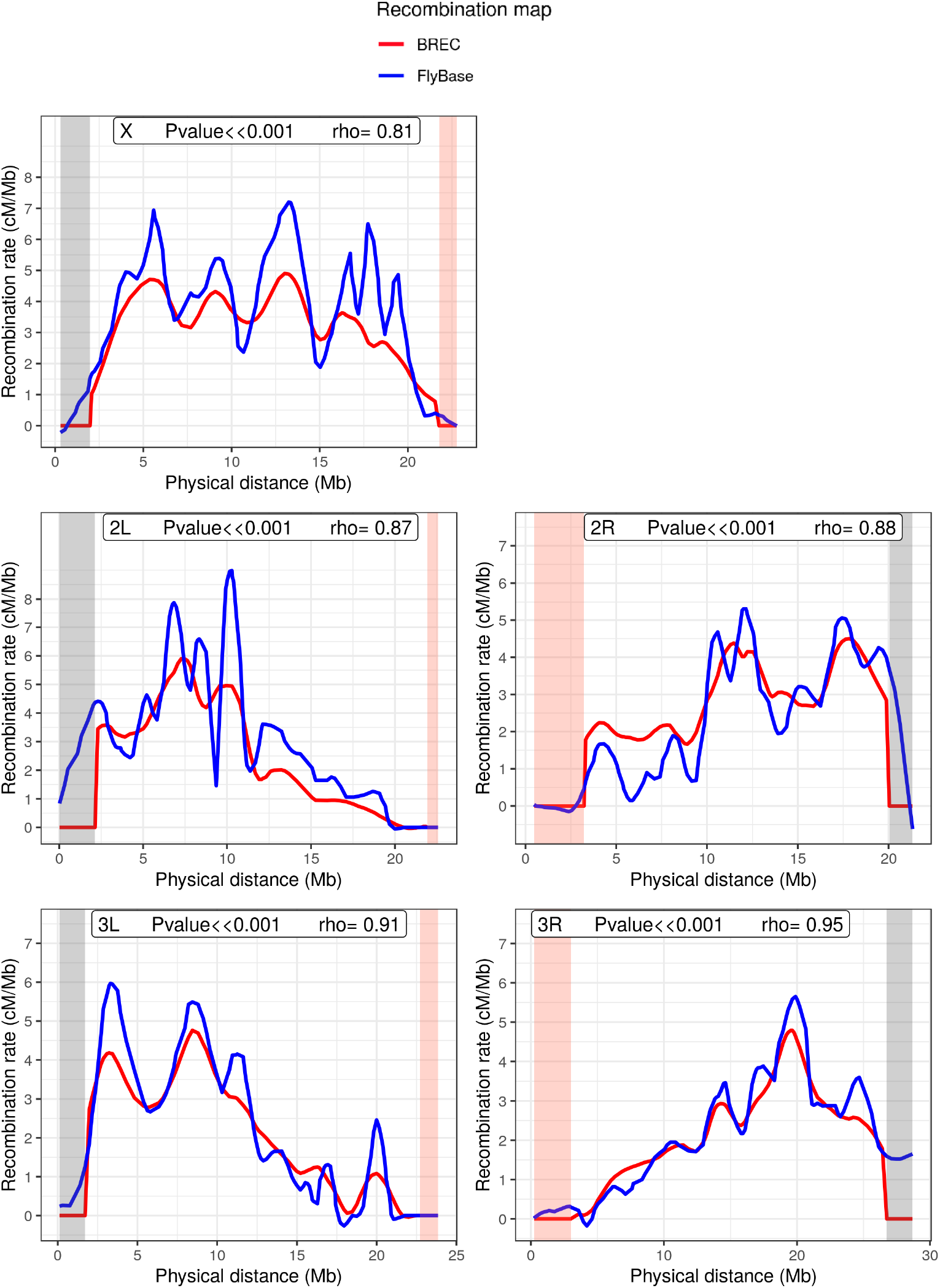
Comparison of BREC *vs*. FlyBase recombination rate recombination rates along the five chromosomal arms (X, 2L, 2R, 3L, 3R) of *D. melanogaster* Release 5. Both recombination maps are obtained using the same regression model: Loess with span 15%. The HCB defined by BREC are represented in red and the reference data are in blue. Heterochromatin regions identified by BREC are highlighted in yellow.

### BREC is non-genome-specific

NGS, High Throughput Sequencing (HTS) technologies and numerous further computational advances are increasingly providing genetic and physical maps with more and more accessible markers along the centromeric regions. Such shift on the availability of data of poorly accessible genomic regions is a huge opportunity to shift our knowledge of the biology and dynamics of heterochromatin DNA sequences as Transposable Elements (TEs) for example. Therefore, BREC is not identifying centromeric gaps as centromeric regions as it might seem, instead, it is targeting centromeric as well as telomeric boundaries identification no matter of the presence or absence of markers neither of their density or distribution variations across such complicated genomic regions (see Fig S12). Given BREC is non-genome-specific, applying HCB identification on various genomes allows to widen the experimental design and to test more thoroughly how BREC responds to different *data scenarios*. Despite the several challenges due to data quality issues and following a data-driven approach, BREC is a non-genome-specific tool that aims to help tackling biological questions.

### Easy, fast and accessible tool *via* an R-package and a Shiny app

BREC is an R-package entirely developed in R programming language. Current version of the package and documentation are available on the GitHub repository: https://github.com/ymansour21/BREC

In addition to the interactive visual results provided by BREC, the package comes with a web-based Graphical User Interface (GUI) build using the shiny and shinydashboard libraries. The intuitive GUI makes it a lot easier to use BREC without struggling with the command line (see screenshots in Figs 3 and S13).

As for the speed aspect, BREC is quite fast when executing the main functions. We reported the running time for *D. melanogaster R5* and *S. lycopersicum* in Tables S1 and S2, respectively (plotting excluded). Nevertheless, when running BREC *via* the Shiny application, and due to the interactive plots displayed, it takes longer because of the plotly rendering. Still, it depends on the size of the genetic and physical maps used, as well as the markers density, as slightly appears in the same tables. The results presented from other species (see Fig S12) highlight better this dependence.

**Fig 3.**
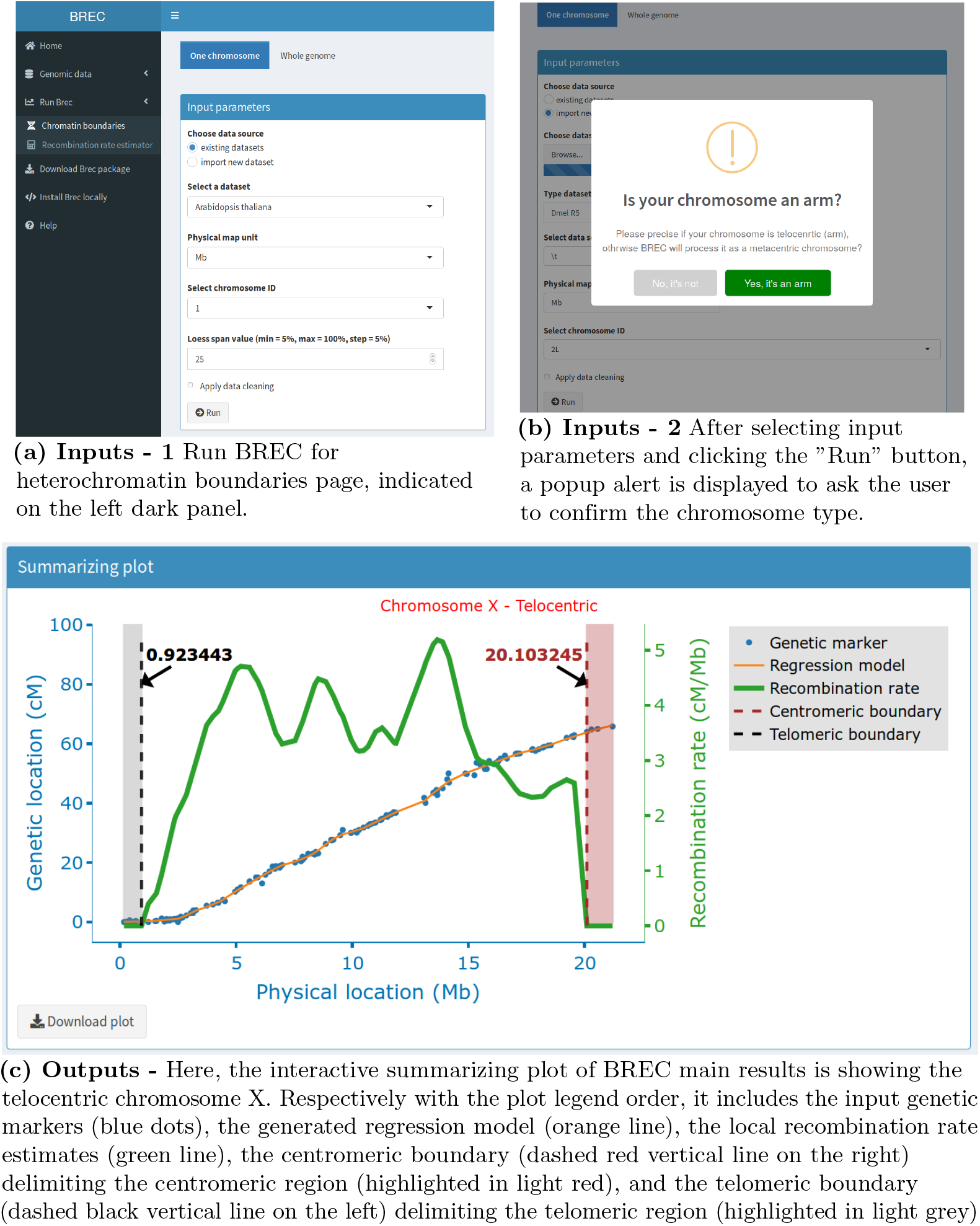
Screenshots of BREC web application - Run BREC web page. (3a) and (3b) show the inputs interface. (3c) shows the output of running BREC on the specified inputs, represented with an interactive web-based plot as a result.

## Discussion

The main two results of BREC are the eu-heterochromatin boundaries and the local recombination rate estimates (see Figs S4; 2).

The HCB algorithm, which identifies the location of centromeric and telomeric regions on the physical map, relies on the regression model obtained from correlation the physical distance and the genetic distance of each marker. Then, the goodness-of-fit measure, the R-squared, is used to obtain a curve upon which the transition between euchromatin and heterochromatin is detectable.

On the other hand, the recombination rate algorithm, which estimates local recombination rates, returns the first derivative of the previous regression model as the recombination rates, then, resets the derivative values to zero along the heterochromatin regions identified (see Fig 1).

We validated the BREC method with a reference dataset known to be of high quality: *D. melanogaster*. While two distinct approaches were respectively implemented for the detection of telomeric and the centromeric regions, our results show a similar high resolution (see Table 1 and Fig S4). Then we analysed BREC’s robustness using simulations of a progressive data degradation (see Figs S8; S10). Even if BREC is sensitive to the markers distribution and thus the local marker density, it can correctly handle a low global marker density. For *D. melanogaster* genome, a density of 5 markers/Mb seems to be sufficient to detect precisely the HCB.

We also validated BREC using the domesticated tomato *S. lycopersicum* dataset (see Table S3 and Fig S7). At first glance, one might ask: why validating with this species when the results do not seem really congruent? In fact, we have decided to investigate this genome as it provides a more insightful understanding of the data-driven aspect of BREC and how data quality strongly impacts the heterochromatin identification algorithm. Variations in the local density of markers in this genome are particularly associated with the relatively large plateaued centromeric region representing more than 50% of the chromosome’s length. Such *data scenario* is quite different compared to what we previously reported on the *D. melanogaster* chromosomal arms. This is partially the reason for which we chose this genome for testing BREC limits. While analysing the experiments more closely, we found that BREC processes some of the chromosomes as presenting a centromeric gap, while that is not actually the case. Thus, we forced the HCB algorithm to automatically apply the *with-no-centromeric-gap-algorithm*, then, we were inspired to implement this option into the GUI in order to give the users the ability to take advantage of their *a priori* knowledge and by consequence to use BREC more efficiently. Meanwhile, we are considering how to make BREC completely automated regarding this point for an updated version later on. In addition, the reference heterochromatin results we used for the BREC validation are in fact rather an approximate than an exact indicator. The reference physical used correspond to the first and last markers tagged as “heterochromatin” on the spreadsheet file published by the Tomato Genome Consortium authors in [32]. However, we hesitated before validating BREC results with these approximate reference values due to the redundant existence of markers tagged as “euchromatin” directly before or after these reference positions. Unfortunately, we were not able to validate telomeric regions since the reference values were not available. As a result, we are convinced that BREC is approximating well enough in the face of all the disrupting factors mentioned above.

On the other hand, the ambition of this method is to escape species-dependence, which means it is conceived to be applicable to a various range of genomes. To test that, we thus also launched BREC on genomic data from different species (the house mouse’s chromosome 4, roundworm’s chromosome 3 and the chromosome 1 of zebrafish). Experiments on these whole genomes showed that BREC works as expected and identifies chromosome types in 95% of cases (see Fig S12).

One can assume, with the exponential increase of genomics resources associated with the revolution of the sequencing technologies, that more and more fine-scale genetic maps will be available. Therefore, BREC has quite the potential to widen the horizon of deployment of data science in the service of genome biology and evolution. It will be important to develop a dedicated database to store all these data.

BREC package and design offer numerous advantageous (see Table S5) compared to similar existing tools [21, 22]. Thus, we believe our new computational solution will allow a large set of scientific questions, such as the ones raised by the authors of [5, 34], to be addressed more confidently, considering model as well as non-model organisms, and with various perspectives.

## Conclusion

We designed a user-friendly tool called BREC that analyses genomes on the chromosome scale, from the recombination point-of-view. BREC is a rapid and reliable method designed to determine euchromatin-heterochromatin frontiers on chromosomal arms or whole chromosomes (resp. telocentric or metacentric chromosomes). BREC also uses its heterochromatin boundary results to improve the recombination rate estimates along the chromosomes.

Whole genome version of BREC is a work in progress. Its will allow to run BREC on all the chromosomes of the genome of interest at ones. This version will also present the identified heterochromatin regions on chromosome ideograms. As short-term perspectives for this work, we may consider extending the robustness tests to other datasets with high quality and mandatory information (*e.g*. boundaries identified with cytological method, high quality maps). Retrieving such datasets seems to become less and less difficult. As well, we may improve the determination of boundaries with a finer analysis around them, for instance using an iterative multi-scale algorithm. Finally, we will be happy to take into account users feedback and improve the ergonomy and usability of the tool. As mid-term perspectives, we underline that BREC could integrate other algorithms aiming to provide further analysis options such as the comparison of heterochromatin regions between closely related species.

## Acknowledgments

This work has been supported by the Algerian Ministry of Higher Education and Scientific Research as part of the Algerian Excellency Scholarship Program AVERROES. This work was also funded by the Junior Research Group (ERJ 2018) grant from Labex CeMEB (ANR). Our thanks go to the members of the “Phylogeny and Molecular Evolution” from the Institute of Sciences and Evolution of Montpellier (ISEM) and the “Methods and Algorithms in Bioinformatics” from the Laboratory of Computer Science, Robotics and Microelectronics of Montpellier (LIRMM).

## Supporting materials

### Materials and Methods

#### Validation data

The only input dataset to provide for BREC is genetic and physical maps one or several chromosomes. A simple CSV file with at least two columns for both maps is valid. If the dataset is for more than one chromosome or for the whole genome, a third column, with the chromosome identifier, is required.

Our results have been validated using the Release 5 of the fruit fly *D. melanogaster* [35, 36] genome as well as the domesticated tomato *Solanum lycopersicum* genome (version SL3.0).

We also tested BREC using other datasets of different species: house mouse (*Mus musculus castaneus*, MGI) chromosome 4 [37], roundworm (*Caenorhabditis elegans*, ws170) chromosome 3 [38], zebrafish (*Danio rerio*, Zv6) chromosome 1 [39], respectively (see Fig S12), as samples from the multi-genome dataset included within BREC (see Table S4).

##### Fruit fly genome *D.melanogaster*

Physical and genetic maps are available for download from the FlyBase website (http://flybase.org/; Release 5) [25]. This genome is represented here with five chromosomal arms : 2L, 2R, 3L, 3R and X (see Table S1), for a total of 618 markers, 114.59Mb of physical map and 249.5cM of genetic map. This dataset is manually curated and is already clean from outliers. Therefore, the cleaning step offered within BREC was skipped.

##### Tomato genome *S. lycopersicum*

Domesticated tomato with 12 chromosomes has a genome size of approximately 900Mb. Based on the latest physical and genetic maps reported by the Tomato Genome Consortium [32], we present both maps content (markers number, markers density, physical map length and genetic map length) for each chromosome in Table S2. For a total of 1957 markers, 752.47Mb of physical map and 1434.49cM of genetic map along the whole genome.

#### Simulated data for quality control testing

We call *data scenarios*, the layout in which the data markers are arranged along the physical map. Various *data scenarios*, for experimentally testing the limits of BREC, have been specifically designed based on *D. melanogaster* chromosomal arms (see Fig S9).

In an attempt to investigate how markers density vary within and between the five chromosomal arms of *D. melanogaster* Release 5 genome, markers density is analyzed in two ways: locally (with 1-Mb bins) and globally (on the whole chromosome). Fig S5 shows the results of this investigation where each little box indicates how many markers are present within each bin of 1Mb size on the physical map, while global markers density per chromosomes is represented by the mean value. Global markers density per chromosomes is also shown in Table S1 where the values are slightly different. This is due to computing markers density in two different ways with respect to the analysis. Table S1, presenting the genomic features of the validation dataset, shows markers density in Column 3, which is simply the result of the division of markers number (in column 2) by the physical map length (in Column 4). For example, in the case of chromosomal arm X, this gives 165/21.22 = 7*.78markers/Mb*. On the other hand, Fig S5, aimed for analysing the variation of local markers density, displays the mean of of all 1-Mb bins densities which is calculated as the sum of local densities divided by the number of bins, and this gives 165/22 = 7*.5markers/Mb*.

The exact same analysis has been conducted on the tomato genome *S. lycopersicum* where the only difference lies is using 5-Mb instead of 1-Mb bins, due to the larger size of its chromosomes (see Fig S6).

#### Validation metrics

The measure we used to evaluate the resolution of BREC’s HCB is called *shift* hereafter. It is defined as the difference between the observed heterochromatin boundary (*observed_HCB*) and the expected one (*expected,_HCB*) in terms of physical distance (in Mb)(see Equation S0).

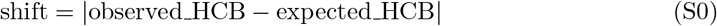

The *shift* value is computed for each heterochromatin boundary independently. Therefore, we observe only two boundaries on a telocentric chromosome (one centromeric and one telomeric) while we observe four boundaries in case of an atelocentric chromosome (two centromeric giving the centromeric region and two telomeric giving each of the two telomeric regions).

The shift measure was introduced not only to validate BREC’s results with the reference equivalents, but also to empirically calibrate the DQC module, where we are mostly interested in the variation of its value with respect to variations of the quality of input data.

#### Implementation and Analysis

The entire BREC project was developed using the R programming language (version 3.6.3 / 2020-02-29) and the RStudio environment (version 1.2.5033). The graphical user interface is build using the shiny and shinydashboard packages. The web-based interactive plots are generated by the plotly package. Data simulations, result analysis, reproducible reports and data visualizations are implemented using a large set of packages such as tidyverse, dplyr, R markdown, Sweave and knitr among others. The complete list of software resources used is available on the online version of BREC package accessible at https://github.com/ymansour21/BREC.

From inside an R environment, the BREC package can be downloaded and installed using the command in the code chunk in Fig S14. In case of installation issues, further documentation is available online on the ReadMe page. If all runs correctly, the BREC shiny application will be launched on your default internet browser (see Shiny interface screenshots in Fig S13 and description of the build-in dataset as well as GUI elements in Supplementary materials).

All BREC experiments have been carried out using a personal computer with the following specs:

- Processor: Intel^®^ Core™ i7-7820HQ CPU @ 2.90GHz x 8
- Memory: 32Mo
- Hard disc: 512Go SSD
- Graphics: NV117 / Mesa Intel^®^ HD Graphics 630 (KBL GT2)
- Operating system: 64-bit Ubuntu 20.04 LTS

### Description of main components of the Shiny app

#### Build-in dataset

Users can either run BREC on a dataset of 40 genomes, mainly imported from [40], enriched with two mosquito genomes from [41] and updated with *D. melanogaster* Release 6 from FlyBase [25] (see Table S4), already available within the package, or, load new genomes data according to their own interest.

User-specific genomic data should be provided as inputs within at least a 3-column CSV file format including for each marker: chromosome identifier, genetic distance and physical distance respectively. On the other hand, outputs from BREC running results are mainly represented *via* interactive plots.

#### GUI input options

The BREC shiny interface provides the user with a set of options to select as parameters for a given dataset (see figure 3a). These options are mainly necessary in case the user works on his/her own dataset and this way the appropriate parameters would be available to choose from. First, a tab to specify the running mode (one chromosome). Then, a radio button group to choose the dataset source (existing within BREC or importing new dataset). For the existing datasets case, there is a drop-down scrolling list to select one of the available genomes (over 40 options), a second one for the corresponding physical map unit (Mb or pb) and a third one for the chromosome ID (available based on the dataset and not the genome biologically speaking). While for the import new dataset case, three more objects are added (see Fig 3b); a fileInput to select csv data file, a textInput to enter the genome name (optional), and a drop-down scrolling list to select the data separator (comma, semicolon or tab character -set as the default-). As for the Loess regression model, the span parameter is required. It represents the percentage of how many markers to include in the local smoothing process. There is a numericInput object set by default at value 15% with an indication about the range of the span values allowed (min = 5%, max = 100%, step = 5%). The user should keep in mind that the span value actually goes from zero to one, yet, in a matter of simplification, BREC handles the conversion on it’s own. Thus, for example, a value of zero basically means that no markers are used for the local smoothing process by Loess, and so, it will induce a running error. Lastly, there is a checkbox to apply data cleaning if checked. Otherwise, the cleaning step will be skipped. This options could save the user some running time if s/he already have a priori knowledge that a specific genome’s dataset has already been manually curated). The user is then all set to hit the Run button. BREC will start processing the chromosome of interest by identifying its type (telocentric or atelocentric). Since this step is quite difficult to automatically get the correct result, the user might be invited to interfere *via* a popup alert asking for a chromosome type confirmation (see Fig 3b). As shown in Fig (S13a), all available genomes could be accessed from the left-hand panel (in dark grey) and specifically on the tab “Genomic data” where two pages are available: “Download data files” which provides a data table corresponding to the selected genome on a scrolling list along with download buttons, and “Dataset details” displaying a more global overview of the whole build-in aata repository (see Fig S13b). To give a glance at the GUI outputs, Fig 3c shows BREC results displayed within an interactive plot where the user will have the an interesting experience by hovering over the different plot lines and points, visualising markers labels, zooming in and out, saving a snapshot as a PNG image file, and many more available options thanks to the plotly package.

## Supplementary figures

**Fig S1.**
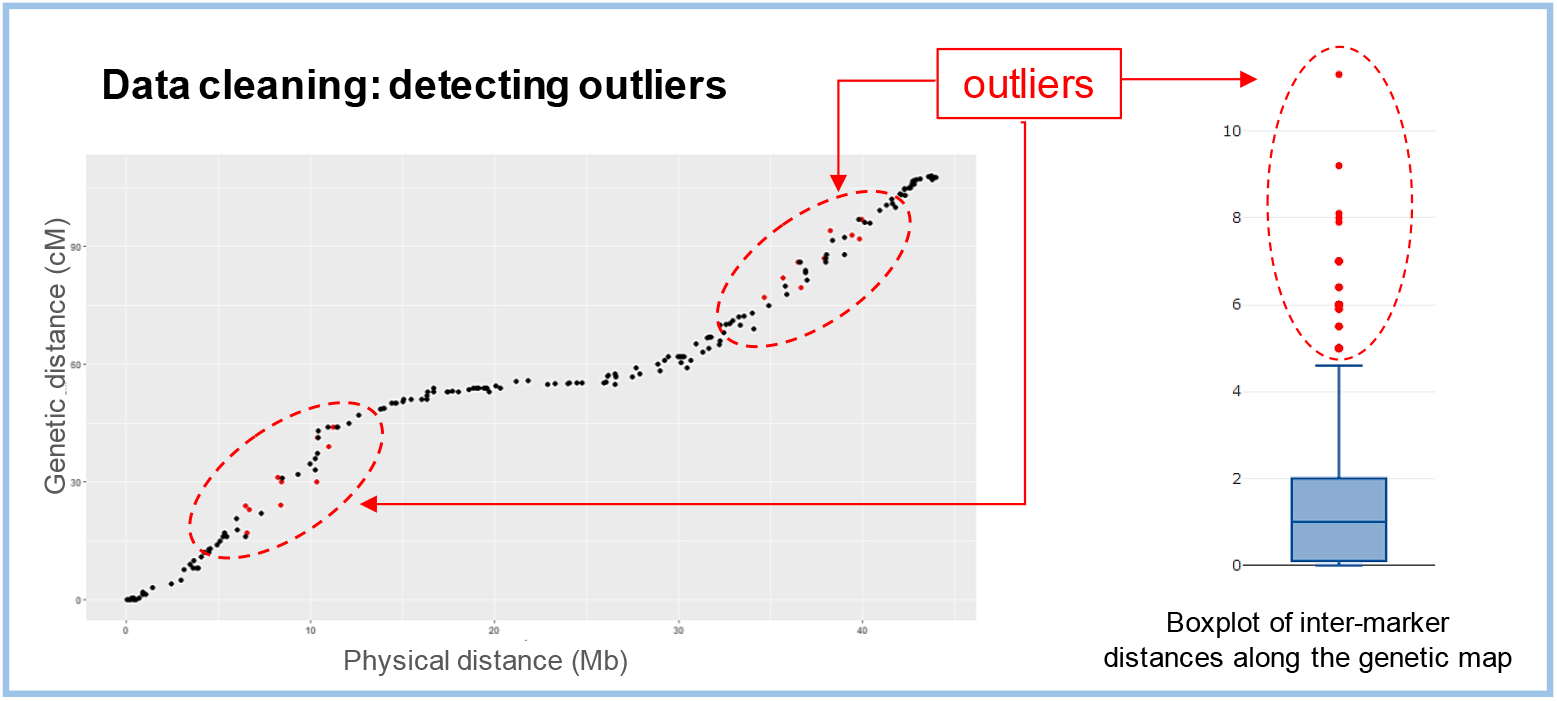
The data cleaning process implemented within BREC. Inter-marker distances (*i.e*. genetic distances between each two consecutive points along the genetic map) are represented using a boxplot in order to identify outliers and give the user the option to remove them. Here is an example showing raw data of a simulated chromosome (left) with the specific markers detected as outliers (red dots circled with red dashed ovals) and the corresponding genetic distances (also in red) on the boxplot (right).

**Fig S2.**
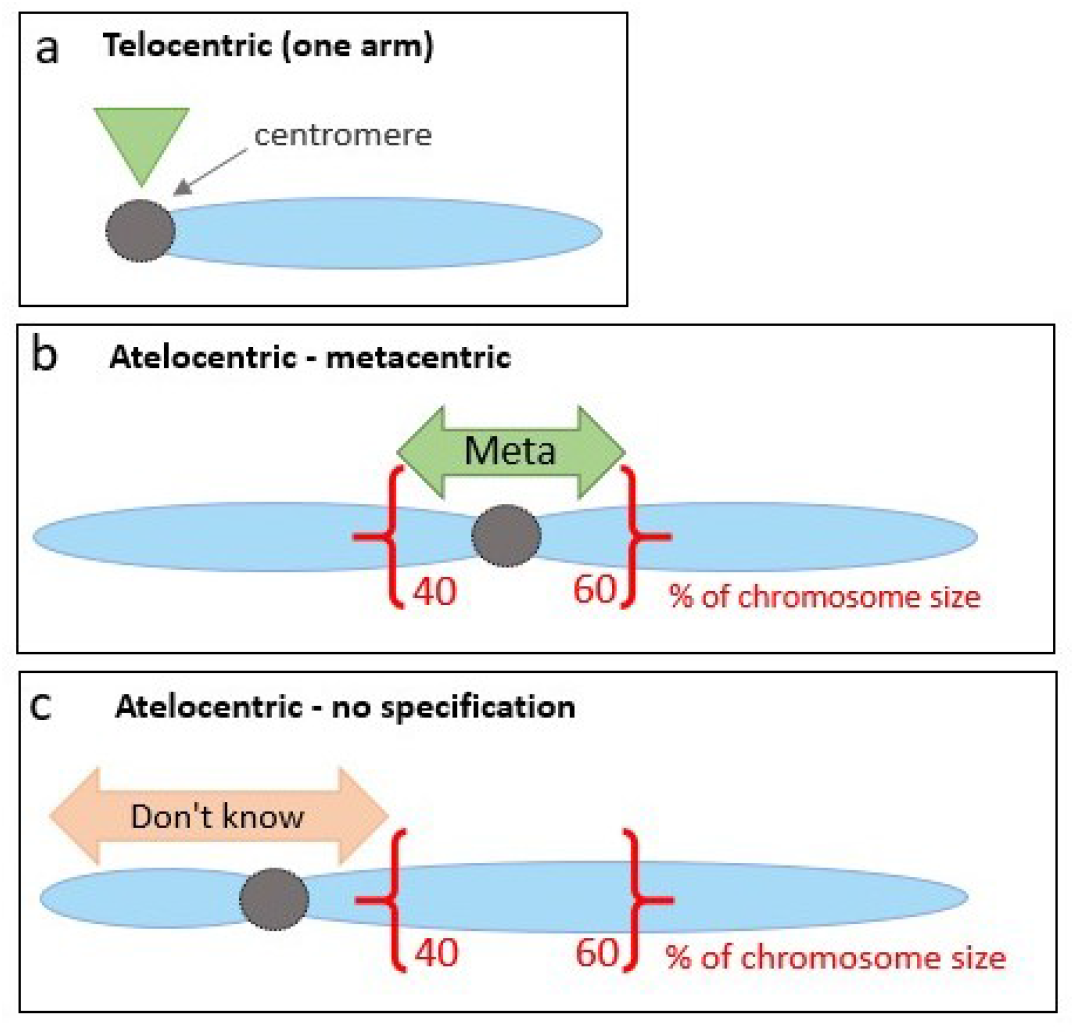
A schematic description of the chromosome type identification process implemented within BREC. (a) Telocentric chromosome type is when the centromere (the grey colored circle) is located on one of the chromosomal arm extremities (indicated with the green upside down triangle). (b) Atelocentric chromosome type -confirmed as metacentric-is when the centromere is located approximately on the middle of the chromosome, here showed within the physical positions 40% and 60% of the chromosome’s size (delimited by the red brackets and indicated with the tag “Meta”). (c) Atelocentric chromosome type -with no specification-is when the centromere is located either inside the first arm (between the beginning of the chromosome and 40% of its size), or inside the second arm (between 60% and the end, indicated with the tag “Don’t know”).

**Fig S3.**
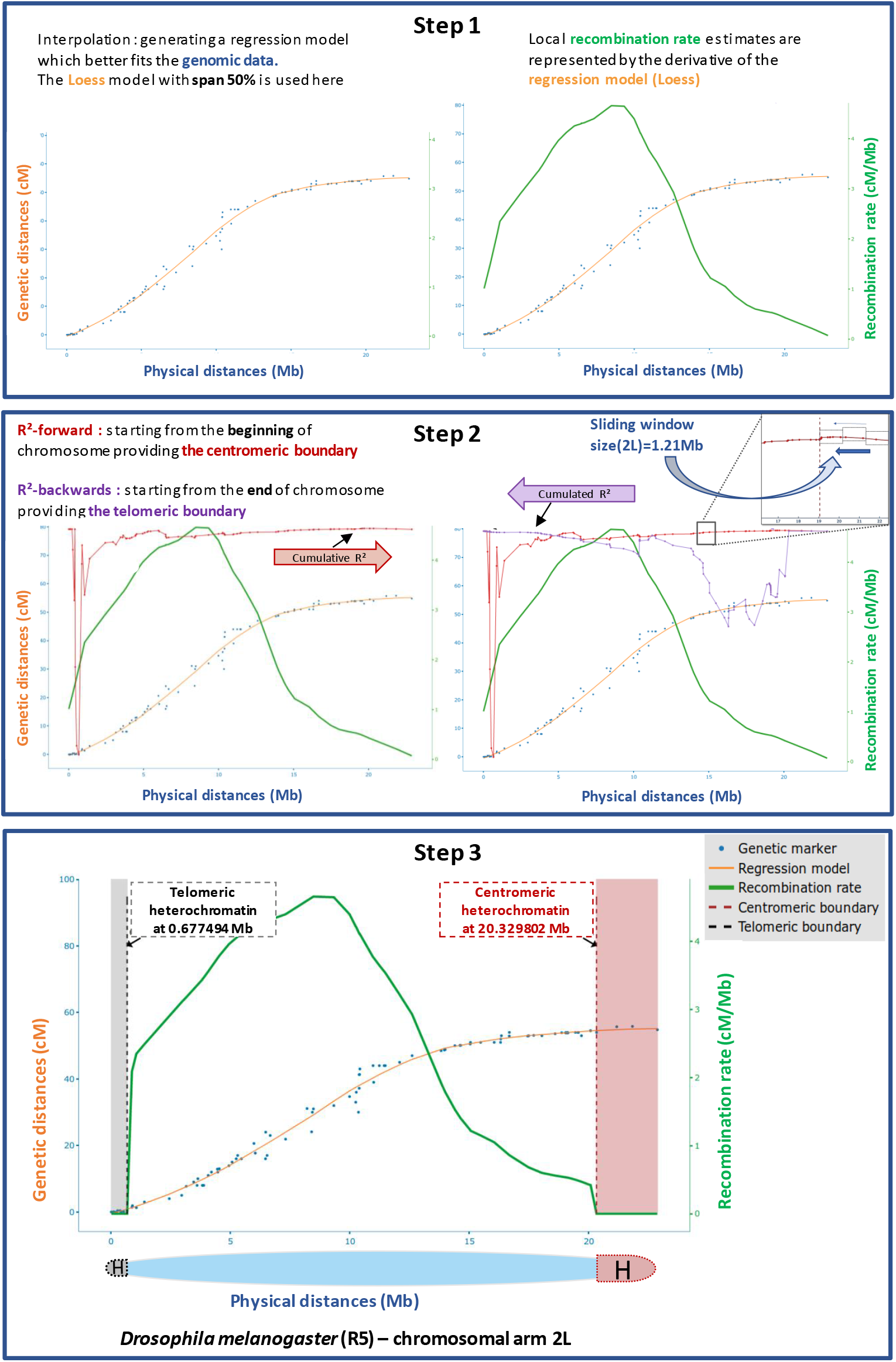
BREC pipeline steps applied on chromosome 2L of *D. melanogaster* Release 5. On each plot, the x-axis represents physical distances (Mb). The left y-axis represents genetic distances (cM) shared between markers (blue data points) and the regression model (orange line). The right y-axis represents recombination rates (cM/Mb) for local estimates (green line). *R*^2^ values, varying between zero and one, are following *R*^2^ – *forward* (red line) and *R*^2^ – *backwards* (purple line). Left telomere and Right centromere (resp. black and purple dashed lines) indicate HCB for the corresponding identified heterochromatin region.

**Fig S4.**
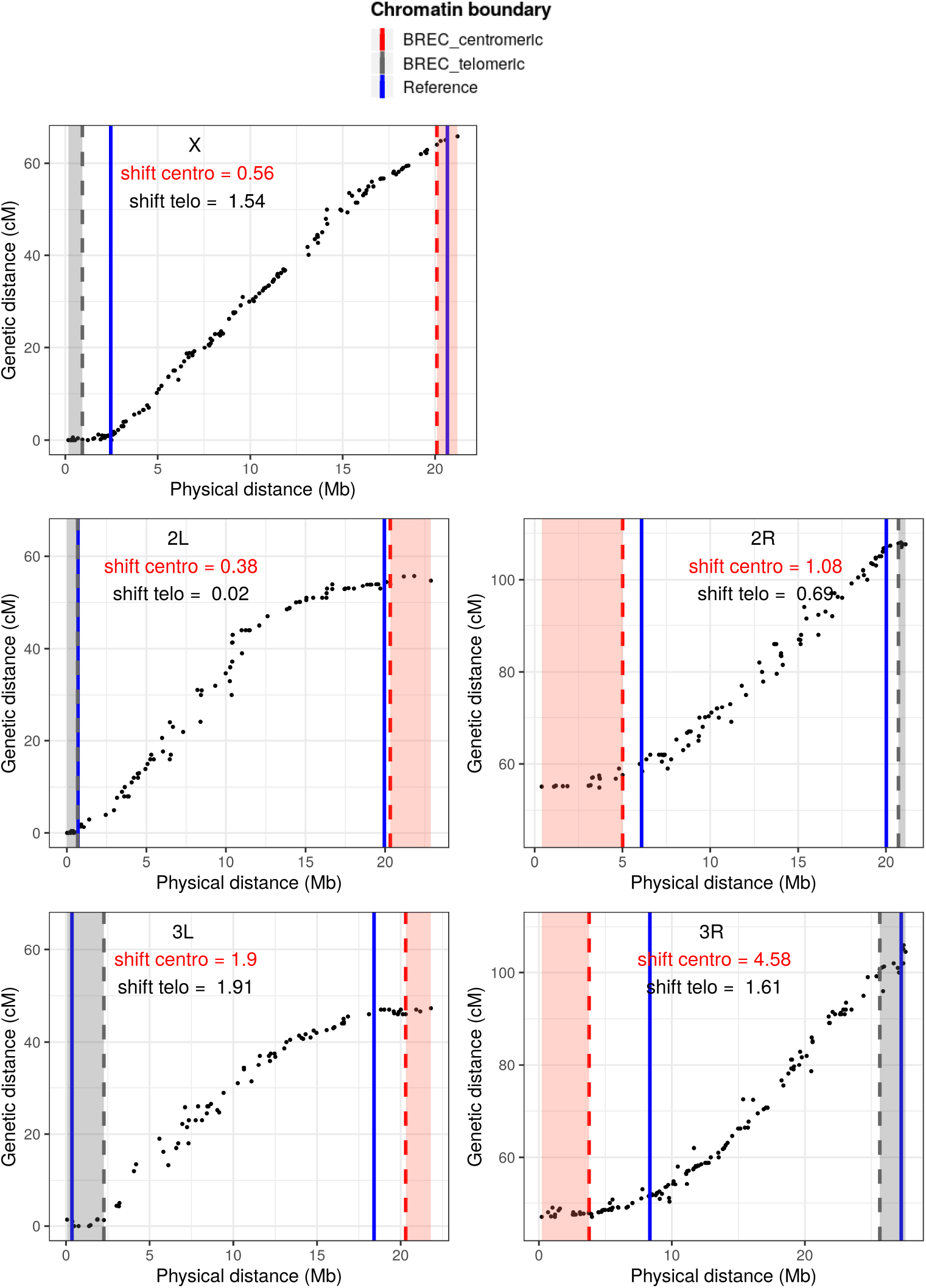
Plots representing results of BREC and reference HCB on the *D. melanogaster* genome. The results are summarized in Table 1. From top to bottom are the five chromosomal arms X, 2L, 2R, 3L, 3R, respectively. Black dots represent genetic markers in ascendant order according to their physical position (in Mb). Vertical lines represent HCB for BREC centromeres (in red dashed line), for BREC telomeres (in grey dashed line) and for the reference (in solid blue line). The heterochromatin regions identified by BREC are highlighted for the centromere (in red) and the telomere (in grey).For each chromosomal arm, two shift values of centromeric and telomeric boundaries are shown under the chromosome identifier.

**Fig S5.**
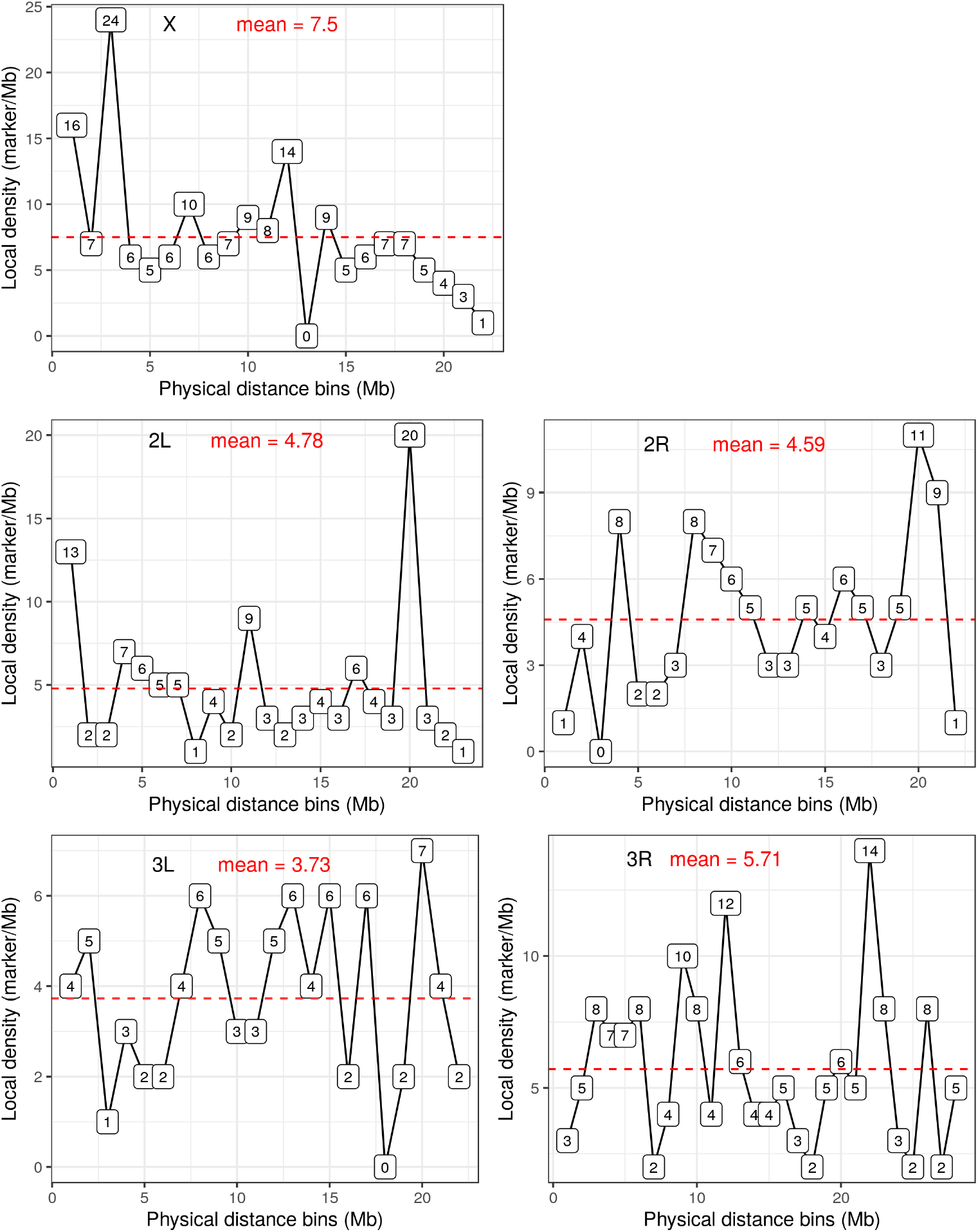
Variations of markers local density per 1-Mb bins along *D. melanogaster* Release 5 chromosomal arms. The red dashed line indicates the mean and represents the global density. Each bin indicates the number of markers it contains. Local density values are represented within the little boxes.

**Fig S6.**
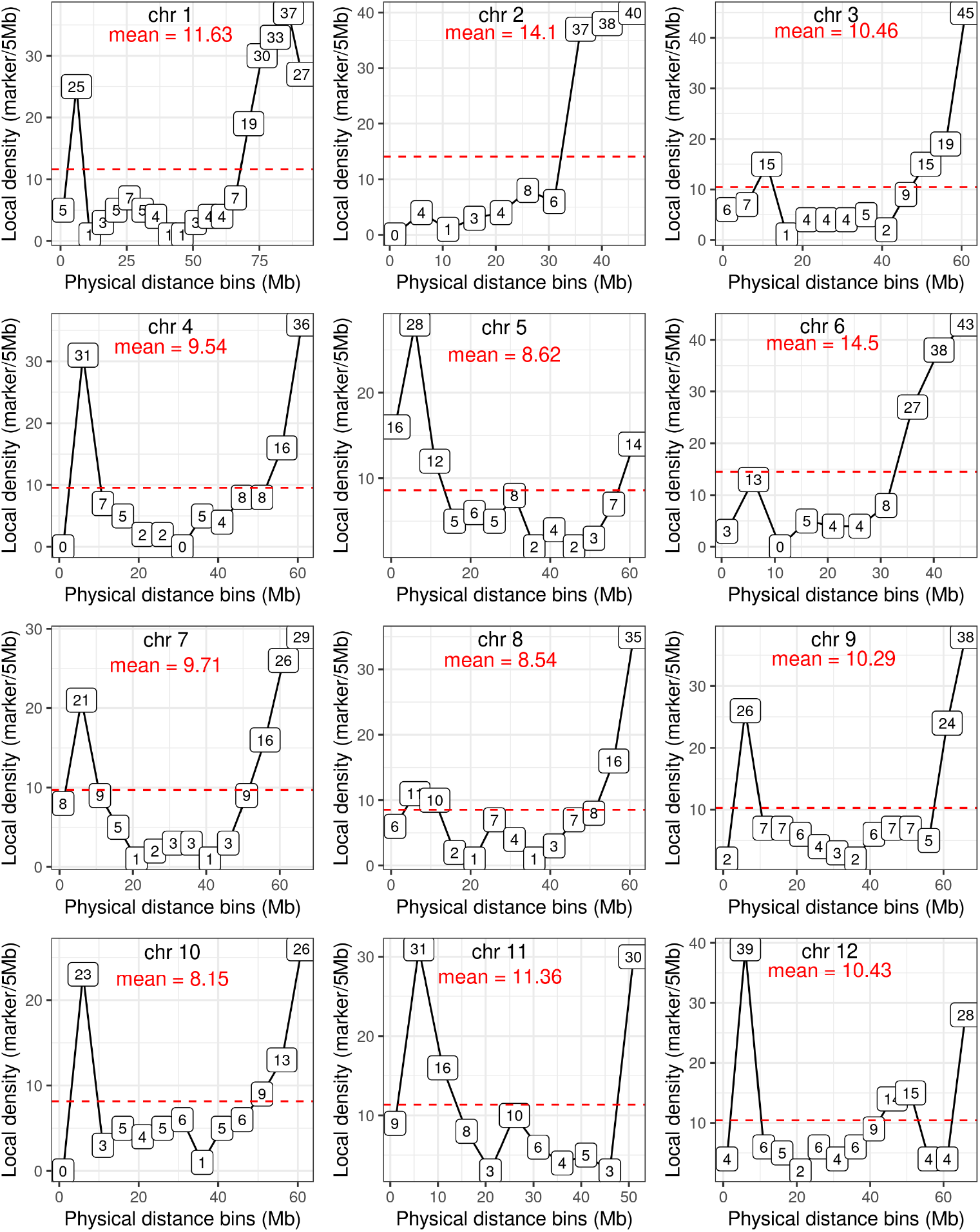
Variations of markers local density per 5-Mb bins along the tomato genome *S. lycopersicum* 12 chromosomes. The red dashed line indicates the mean and represents the global density. Each bin indicates the number of markers it contains. Local density values are represented within the little boxes.

**Fig S7.**
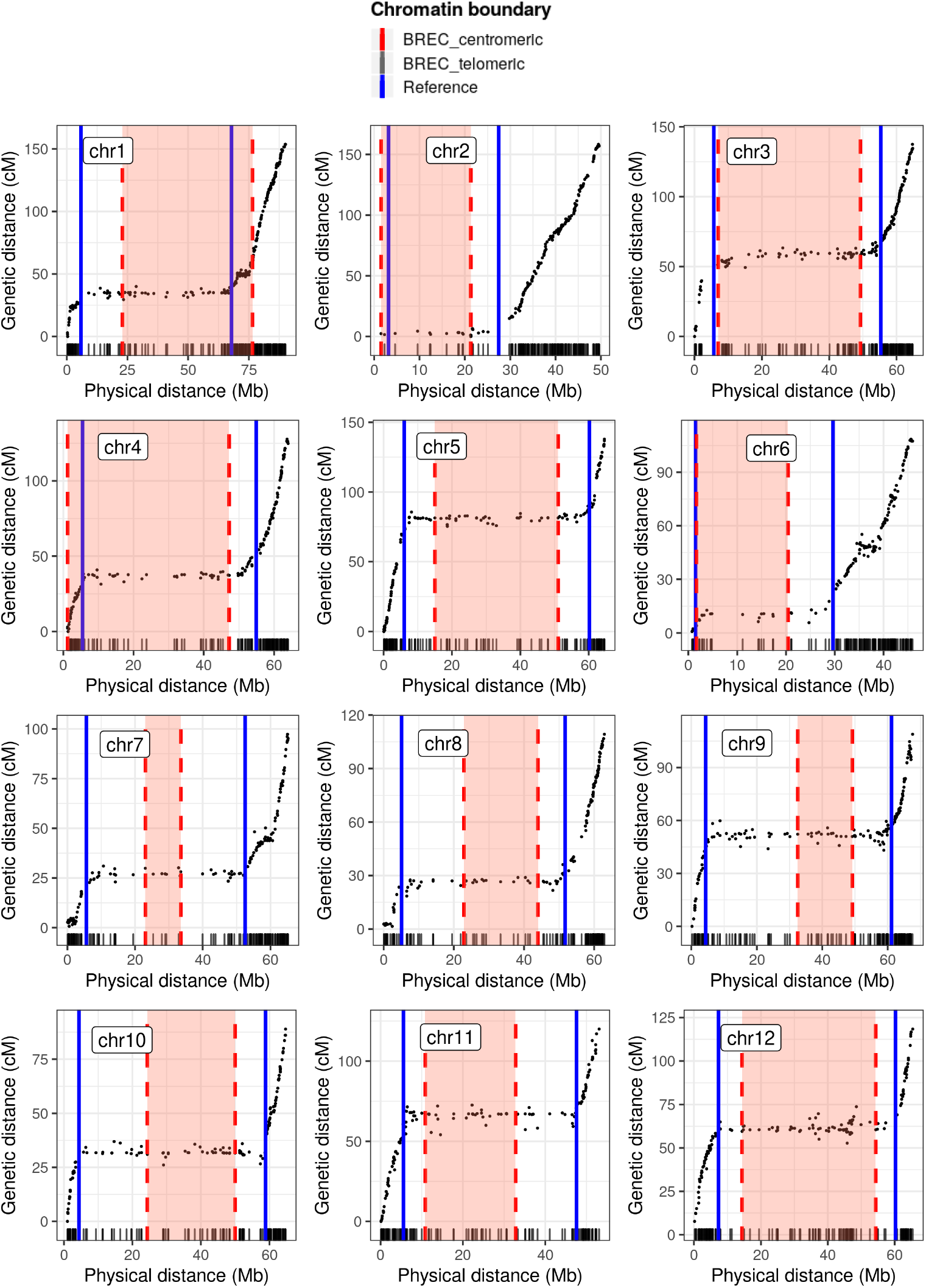
Plots representing results of BREC and reference HCB on the *S. lycopersicum* genome. The results are summarized in Table S3. From top to bottom are the twelve chromosomes 1 to 12, respectively. Black dots represent genetic markers in ascendant order according to their physical position (in Mb). Vertical lines represent HCB for BREC centromeres (in red dashed line), and for the reference (in solid blue line). The heterochromatin regions identified by BREC are highlighted for the centromere (in red). Rug plot on the x-axis represents the markers density according to the physical map.

**Fig S8.**
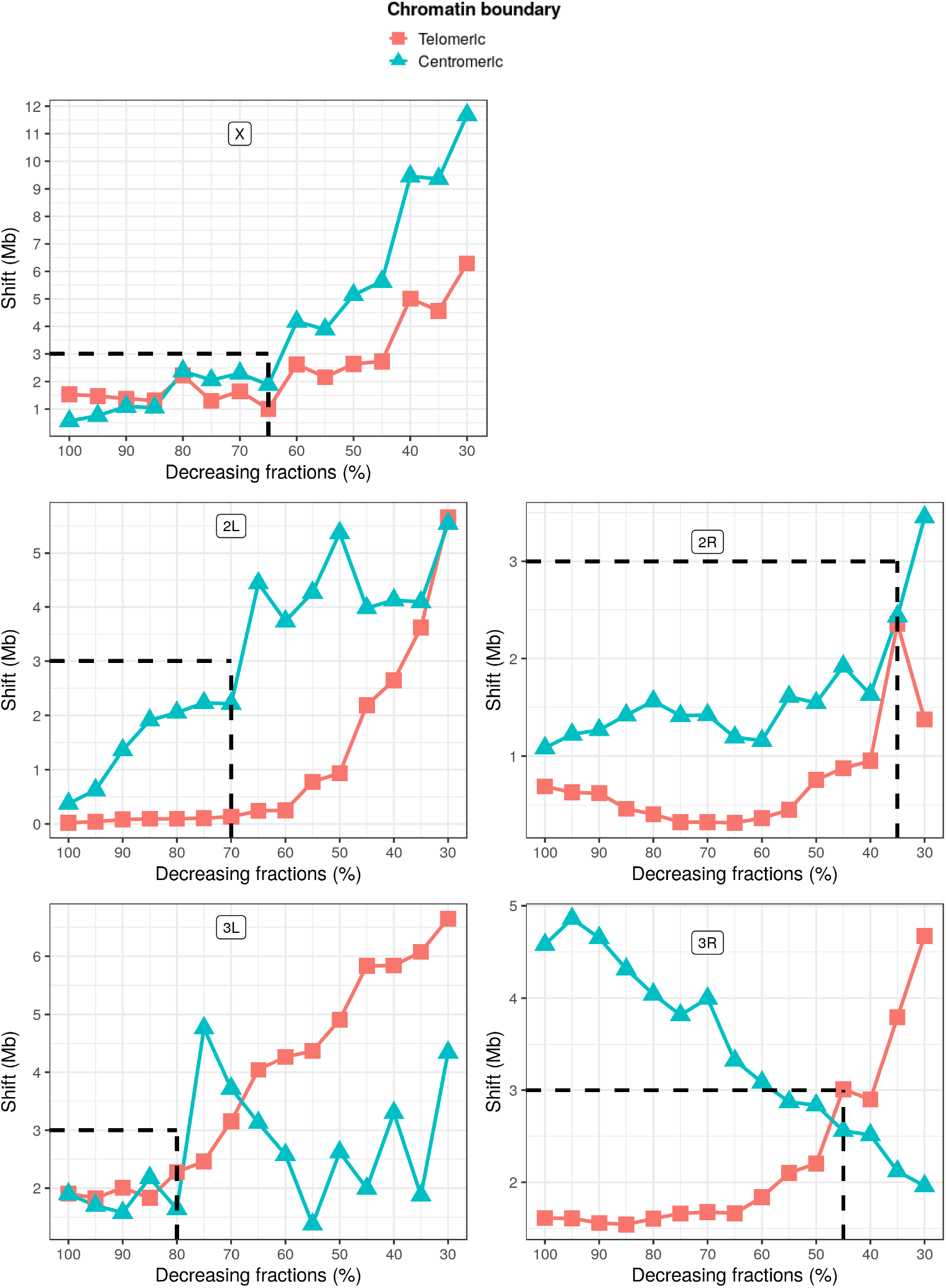
The impact of decreasing markers density on the resolution of BREC’s HCB expressed by the shift value. Here is an overview of the variation of shift values (see Equation S0) for BREC’s HCB compared to reference results for the five *D. melanogaster* chromosomal arms (X, 2L, 2R, 3L, 3R). For each arm, two HCB are shown: squares (in red) for telomeric and triangles (in light blue) for centromeric boundaries. The horizontal dashed line (in black) delimits results smaller than a shift value of 3Mb for all arms while the vertical dashed line (in black) indicates up to which fraction the 3Mb shift is conserved on each chromosomal arm’s simulations. Note that the x axis is reversed, so from left to right it goes from 100% to 30% with a step of −5%at each point. The simulation process is further clarified for one fraction on the chromosomal arm 2L and is illustrated in Fig S10.

**Fig S9.**
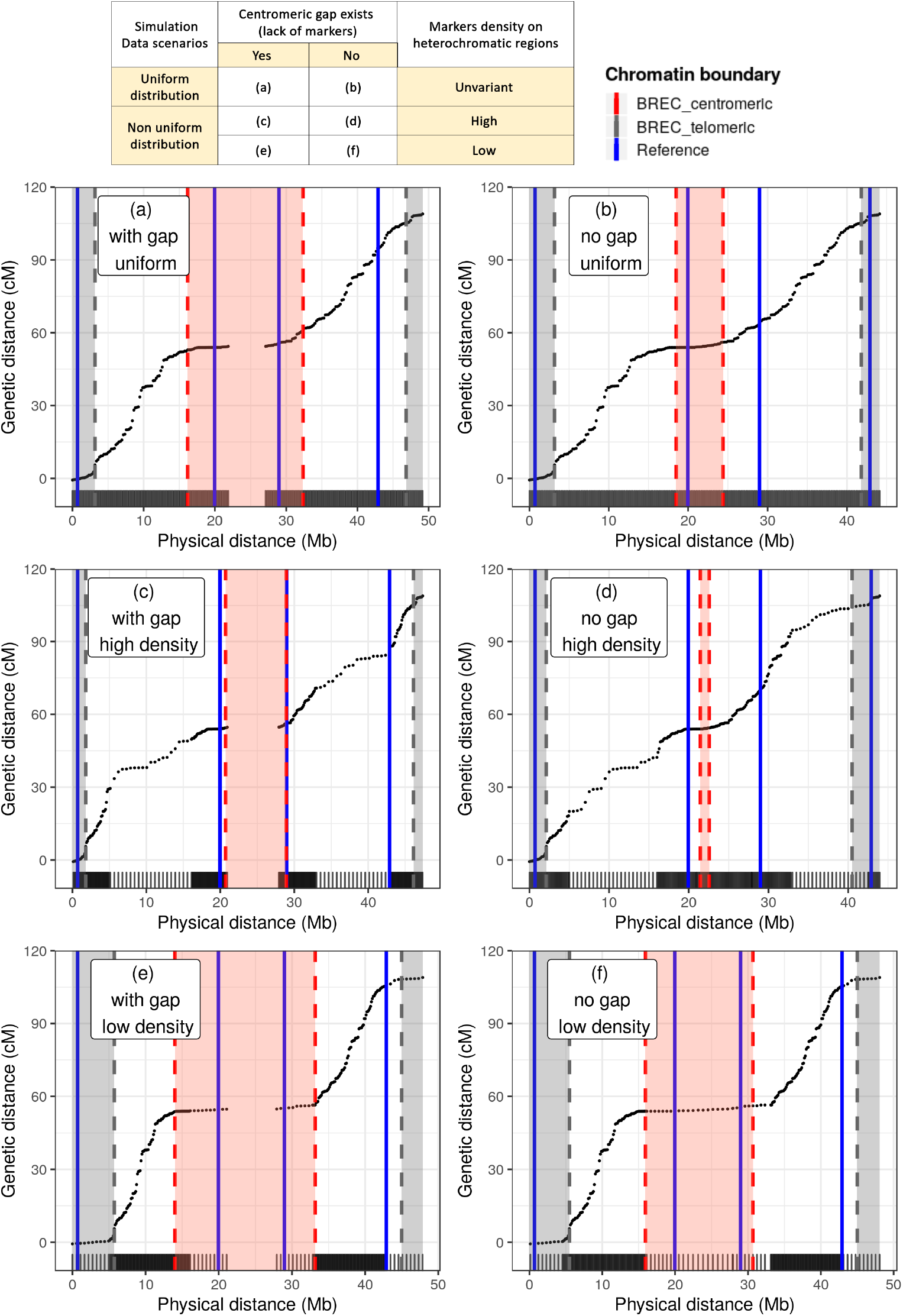
Distribution simulations. BREC results on the simulated chromosomes with different scenarios of markers distribution around heterochromatin regions, as presented in the table (top). Plots (right after) are presenting the corresponding results for each simulation scenario. On the left, (a, c, e) show the cases with the existence of centromeric gap while the ones on the right (b, d, f) show the cases with no centromeric gap. From top to bottom, cases (a) and (b) show a uniform distributions while (c) to (f) are for non uniform distributions. Cases (c) and (d) show a higher density of markers around heterochromatin regions while cases (e) and (f) show a lower density on the same regions. Black dots represent genetic markers. Vertical lines represent HCB for BREC centromeres (in red dashed line), for BREC telomeres (in grey dashed line) and for the reference (in solid blue line). The heterochromatin regions identified by BREC are highlighted for the centromere (in red) and the telomere (in grey). The rug plot, added on the x axis, shows more clearly the variation in markers density as well as the existence or not of the centromeric gap.

**Fig S10.**
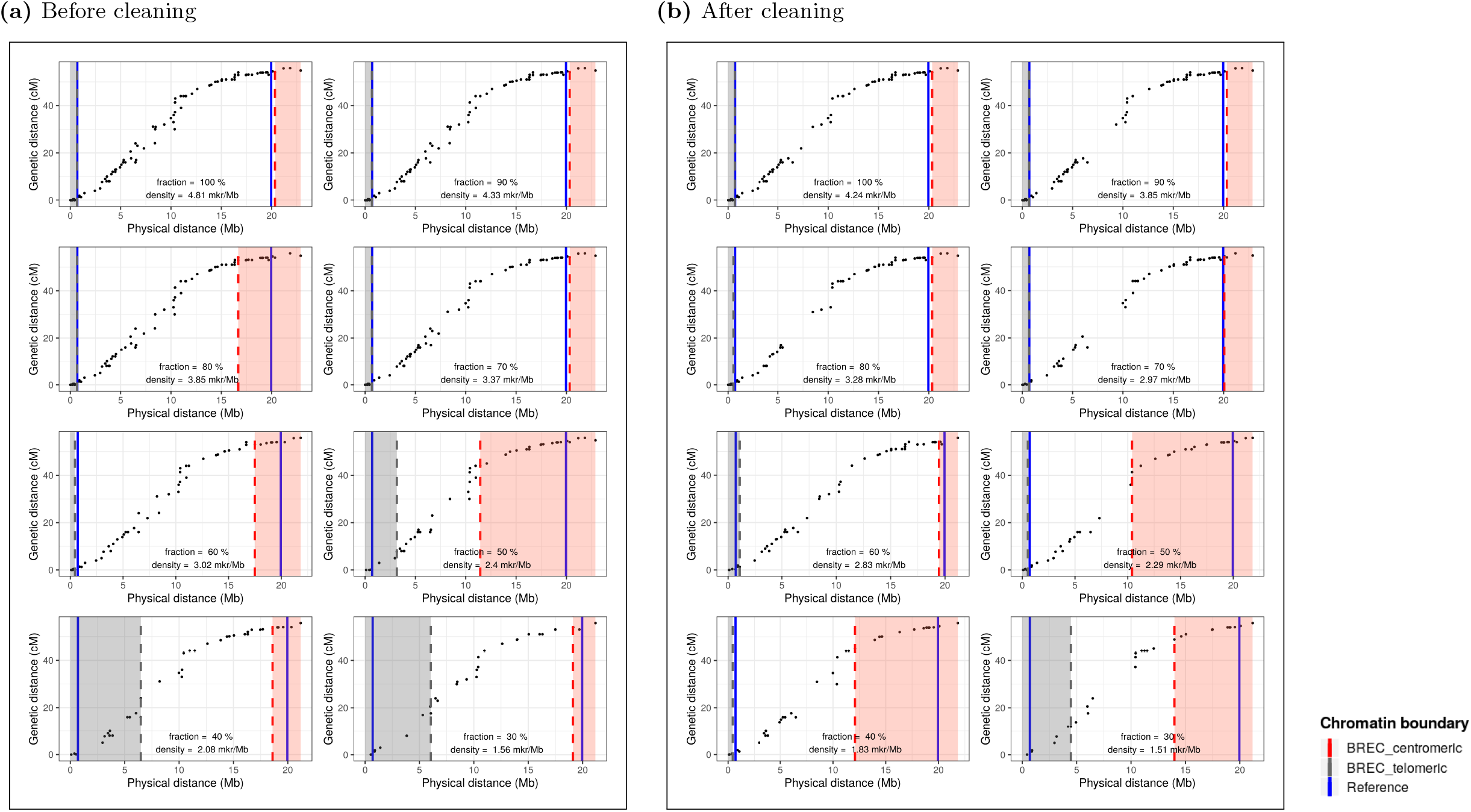
Low density simulations BREC results on the simulated telocentric chromosomes with different density scenarios. Simulating decreasing markers density going from 100% to 30% of the original chromosome 2L (of size 23Mb) of the *D. melanogaster* Release 5 genome. These simulations allow to study the impact of variable markers density on BREC results compared to reference HCB. (a) on the left is before and (b) on the right is after the cleaning step. These simulations have been conducted on each of the five chromosomes (X, 2L, 2R, 3L, 3R) 30 times where the mean shift value is reported in Fig S8. Black dots represent genetic markers. Vertical lines represent HCB for BREC centromeres (in red dashed line), for BREC telomeres (in grey dashed line) and for the reference (in solid blue line). The heterochromatin regions identified by BREC are highlighted for the centromere (in red) and the telomere (in grey). The corresponding fraction and markers density is shown on the top left of each simulation plot.

**Fig S11.**
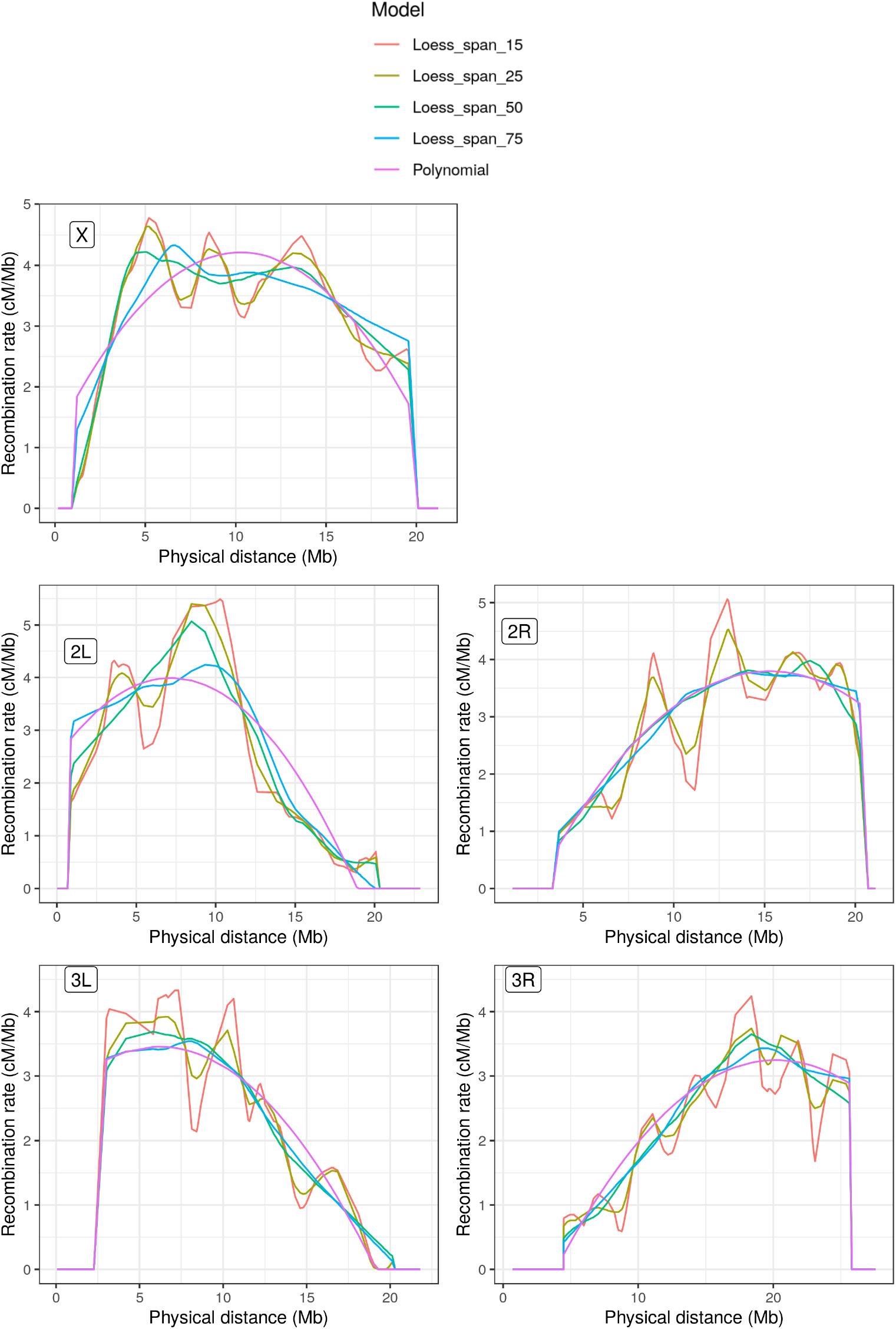
Comparison of regression models for recombination rate estimates along the five chromosomes (X, 2L, 2R, 3L, 3R) of *D. melanogaster* Release 5. Regression models used here are Loess with span values, 15%, 25%, 50%, 75% and third degree polynomial. The HCB defined by BREC remain unchanged and only local recombination rates differ according to the model used to fit the genetic and physical maps. Recombination rate is represented by the derivative of the model. In case of two or more models yielding the same recombination rate estimates on the same physical position, the overlap results in only one curve line. Here, all curves show null recombination rate value on the centromeric and telomeric regions.

**Fig S12.**
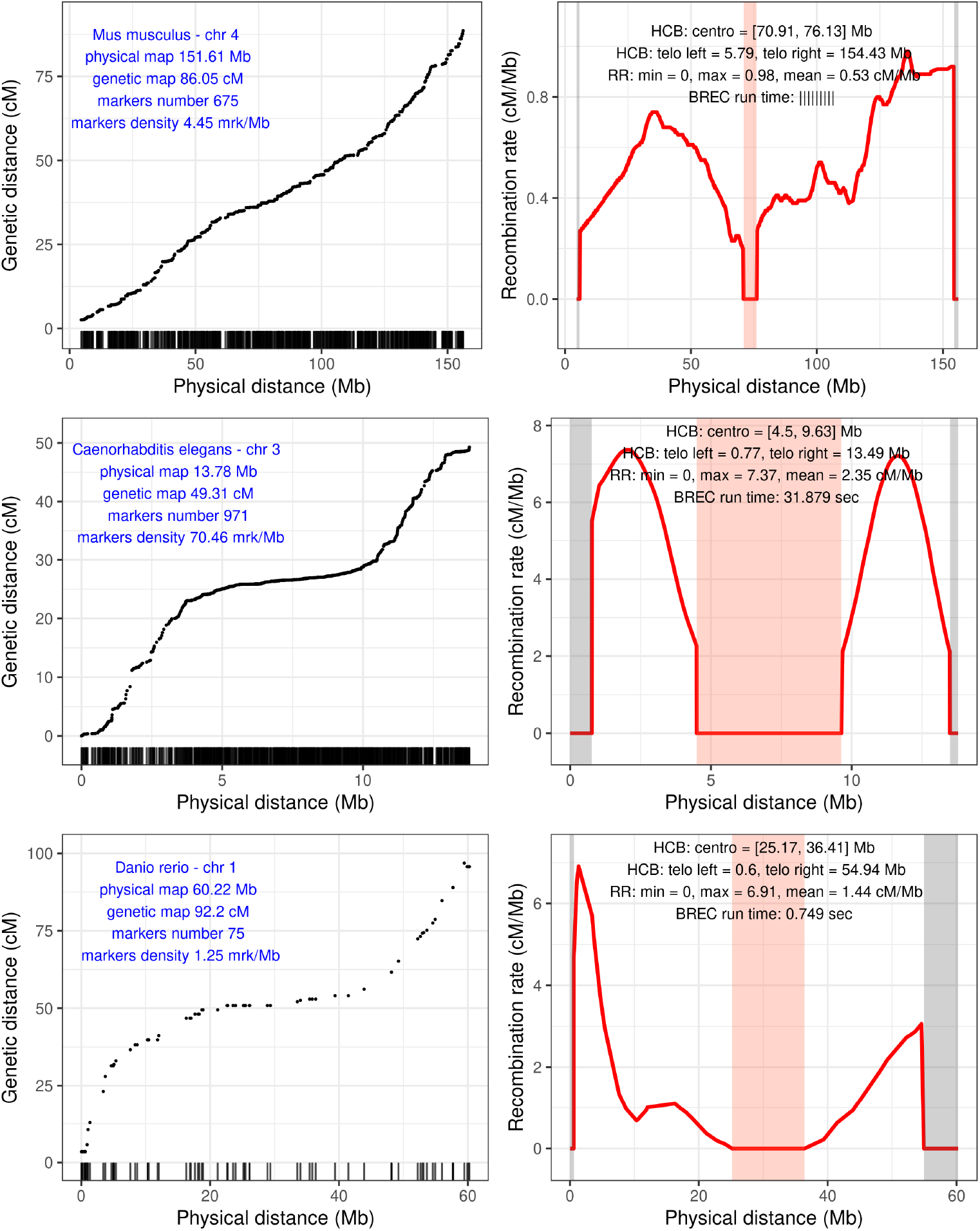
BREC results on different species: from top to bottom are *M. musculus* (house mouse) chromosome 4, *C. elegans* (roundworm) chromosome 3, *D. rereo* (zebrafish) chromosome 1, respectively. For each species, two plots are shown: on the left is the chromosome’s genetic markers (black points), their distribution along the physical map (rug on the x-axis), and reported genomic features (label in blue). On the right is BREC results: HCB for centromeric (red highlight) and telomeric (grey highlight) regions, (RR) local recombination rate estimates (red line), and the running time of BREC’s algorithms to get these results (loading data and plotting are excluded).

**Fig S13.**
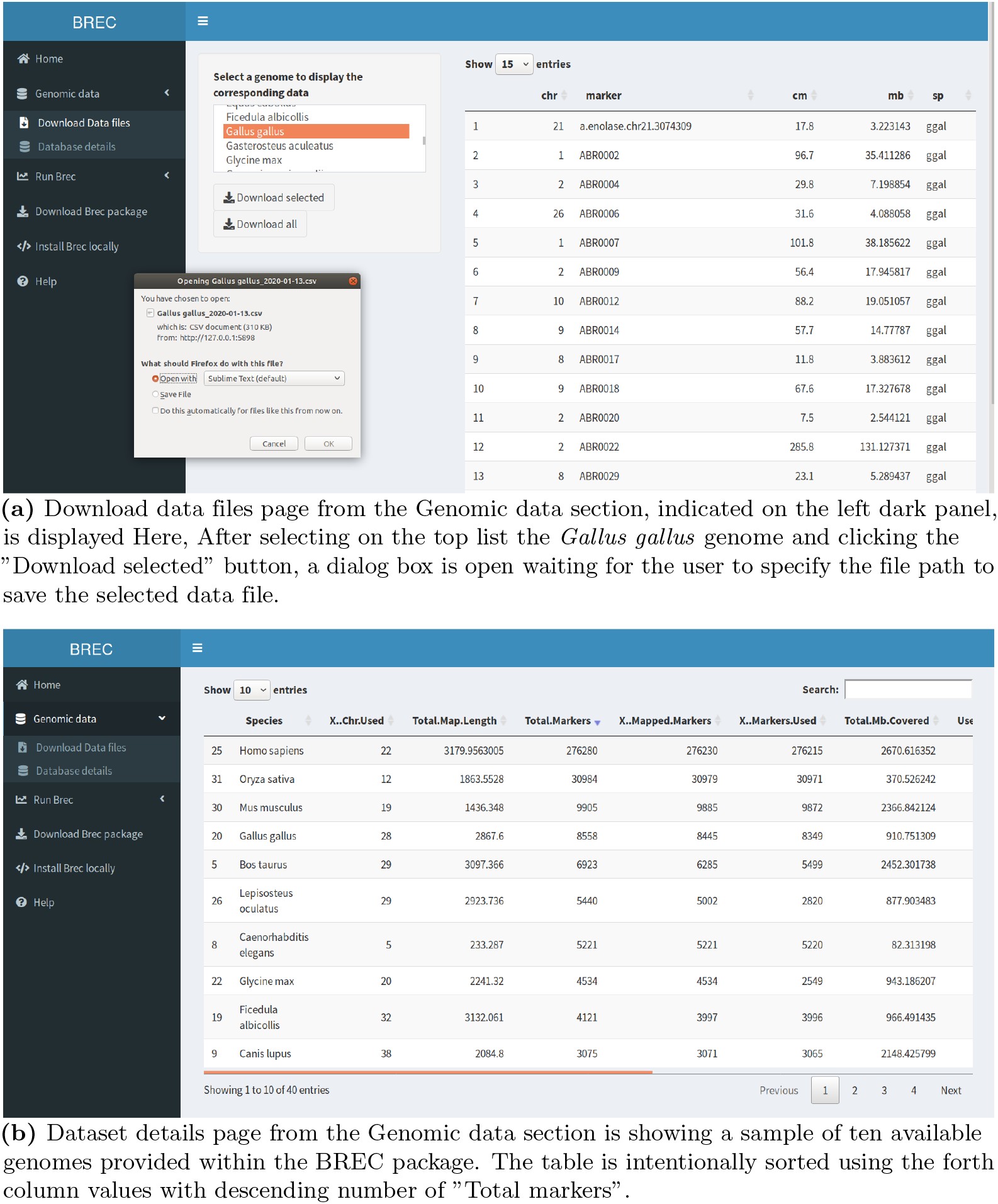
Screenshots of BREC web application - Genomic data web pages. **(a)** Download data files page from the Genomic data section, indicated on the left dark panel, is displayed Here, After selecting on the top list the *Gallus gallus* genome and clicking the “Download selected” button, a dialog box is open waiting for the user to specify the file path to save the selected data file. **(b)** Dataset details page from the Genomic data section is showing a sample of ten available genomes provided within the BREC package. The table is intentionally sorted using the forth column values with descending number of “Total markers”.

**Fig S14.**
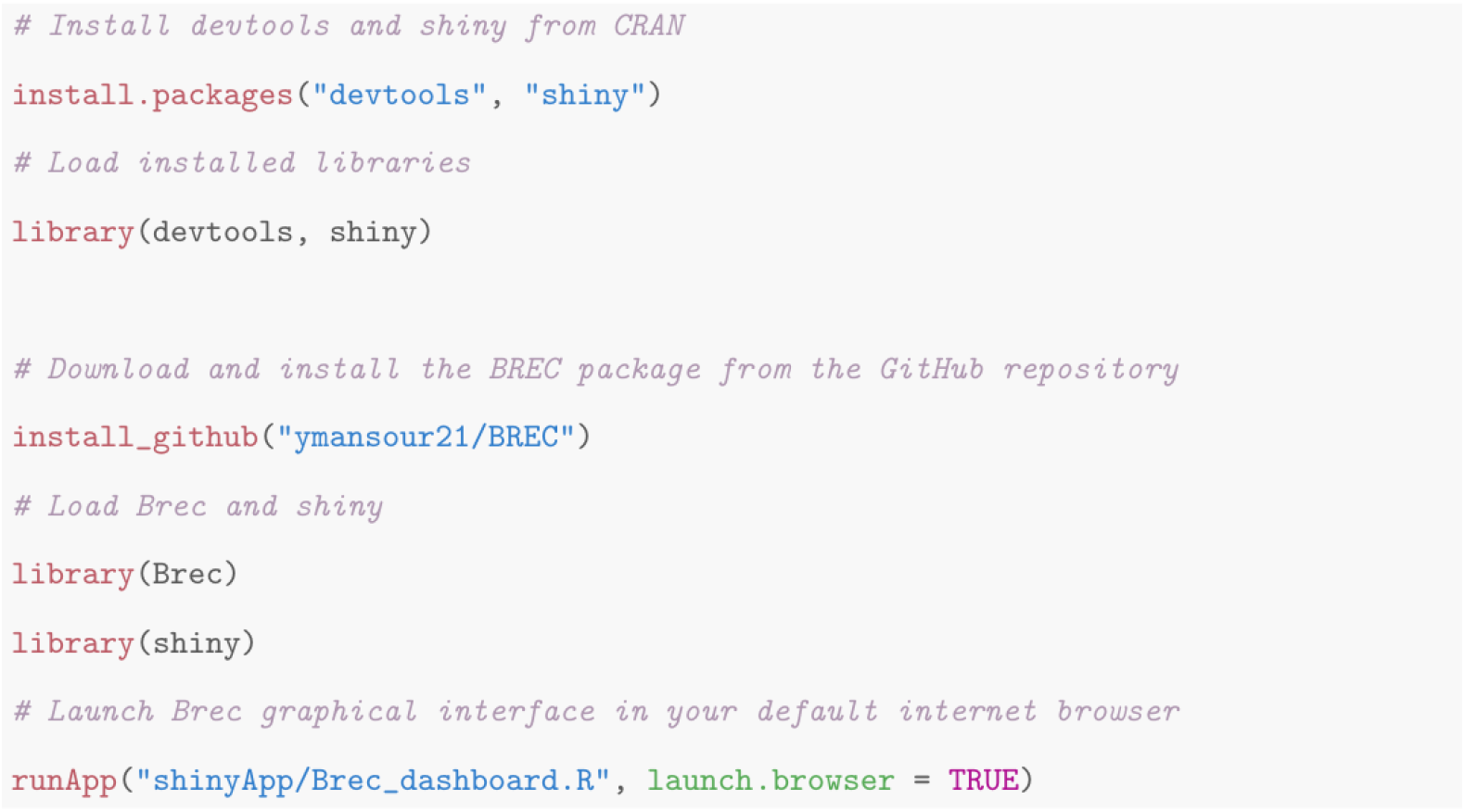
Download, install and launch BREC. Code chunk showing the R commands allowing to download, install and run the BREC shiny application. The entire R package is available with open access on the indicated GitHub repository.

## Supplementary tables

**Table S1.** Genomic features and BREC running time for the *D. melanogaster* Release 5 genome. The first five rows represent chromosomal arms. Columns represent the genome features as follows: (1) the names of chromosomal arms X, 2L, 2R, 3L, and 3R; (2) the markers number included in the study; (3) the markers density (in markers/Mb); and (4) the physical map length (in Mb). The last row summarizes the same features for the whole genome.

| Chromosomal arms | X | 2L | 2R | 3L | 3R | Genome |
| --- | --- | --- | --- | --- | --- | --- |
| Markers number | 165 | 110 | 101 | 82 | 160 | 618 |
| Markers density (marker/Mb) | 7.78 | 4.81 | 4.78 | 3.56 | 5.80 | 5.39 |
| Physical map length (Mb) | 21.22 | 22.88 | 21.12 | 21.81 | 27.57 | 114.59 |
| Genetic map length (cM) | 65.8 | 54.8 | 52.5 | 45.9 | 57.5 | 276.5 |
| BREC run time (sec) | 1.278 | 0.949 | 0.821 | 0.916 | 1.379 | 5.343 |

**Table S2.** Genomic features and BREC running time for the *S. lycopersicum*. The first twelve rows represent chromosomes. Columns represent the genome features as follows: (1) the identifiers of chromosomes 1 to 12; (2) the markers number included in the study; (3) the markers density (in markers/Mb); (4) the physical map length (in Mb); and the elapsed time when running BREC. The last row summarizes the same features for the whole genome.

| Chromosomes | 1 | 2 | 3 | 4 | 5 | 6 | 7 | 8 | 9 | 10 | 11 | 12 | Genome |
| --- | --- | --- | --- | --- | --- | --- | --- | --- | --- | --- | --- | --- | --- |
| Markers number | 232 | 176 | 184 | 160 | 150 | 151 | 145 | 144 | 171 | 148 | 142 | 154 | 1957 |
| Markers density (marker/Mb) | 2.58 | 3.66 | 2.84 | 2.55 | 2.32 | 3.34 | 2.22 | 2.29 | 2.54 | 2.32 | 2.68 | 2.36 | 2.64 |
| Physical map length (Mb) | 89.85 | 48.10 | 64.77 | 62.79 | 64.52 | 45.20 | 65.18 | 62.87 | 67.37 | 63.66 | 52.98 | 65.18 | 752.47 |
| Genetic map length (cM) | 150.72 | 154.58 | 134.52 | 122.64 | 137.91 | 106.63 | 92.48 | 106.63 | 108.90 | 88.92 | 119.99 | 110.72 | 1434.49 |
| BREC run time (sec) | 2.164 | 1.391 | 1.434 | 1.295 | 1.098 | 1.197 | 1.102 | 1.047 | 1.357 | 1.095 | 1.081 | 1.221 | 15.479 |

**Table S3.** Results of BREC and reference HCB on the genome of *S. lycopersicum*. The shift is the absolute value of the distance between the BREC and the reference physical heterochromatin boundary. The first twelve rows represent all chromosomes. Grouped columns present reference, BREC and shift values for the left centromeric boundaries (Columns 2-4), and for the right centromeric boundaries (Columns 4-6). All values are expressed in Megabase (Mb). The red asterisk indicates the largest shift value reported on centromeric and telomeric boundaries separately (see corresponding Fig S7). The last four rows represent some general statistics on the shift value. From top to bottom, they are minimum, maximum, mean, and median respectively. See details on the shift metrics in Section.

| Chromosome | Centromeric left (Mb) |  |  | Centromeric right (Mb) |  |  |
| --- | --- | --- | --- | --- | --- | --- |
|  | Boundaries |  | Shift | Boundaries |  | Shift |
|  | Reference | BREC |  | Reference | BREC |  |
| 1 | 5.78 | 22.88 | 17.09 | 67.80 | 76.48 | 8.68 |
| 2 | 3.15 | 1.51 | 1.64 | 27.43 | 21.31 | 6.12 |
| 3 | 5.75 | 6.98 | 1.23 | 55.34 | 49.28 | 6.06 |
| 4 | 5.48 | 1.21 | 4.27 | 54.92 | 47.21 | 7.72 |
| 5 | 6.02 | 15.03 | 9.01 | 60.23 | 51.04 | 9.19 |
| 6 | 1.50 | 1.68 | 0.19 | 29.62 | 20.42 | 9.20 |
| 7 | 5.62 | 23.05 | 17.43 | 52.51 | 33.52 | 18.98* |
| 8 | 5.10 | 22.87 | 17.77 | 51.73 | 43.96 | 7.77 |
| 9 | 4.38 | 32.51 | 28.12* | 61.16 | 49.16 | 12.00 |
| 10 | 4.40 | 24.37 | 19.97 | 58.83 | 49.92 | 8.91 |
| 11 | 5.56 | 10.86 | 5.29 | 47.57 | 32.77 | 14.80 |
| 12 | 7.27 | 14.34 | 7.07 | 60.27 | 54.33 | 5.94 |
| Min. shift | 0.19 |  |  | 5.94 |  |  |
| Max. shift | 28.12 |  |  | 18.98 |  |  |
| Mean shift | 10.76 |  |  | 9.61 |  |  |
| Median shift | 8.04 |  |  | 8.80 |  |  |

**Table S4.** BREC’s built-in dataset of genomic data. The available genetic and physical maps for 40 species from [40], enriched with two recently assembled mosquito genomes: *Culex pipiens* and *Aedes aegypti* from [41], domesticated tomato *S. lycopersicum* from [32], and *D. melanogaster* Release 6 (update) from FlyBase [25]. The species in red bold text are the one we use in BREC experiments. Since the data collection process is still ongoing, the current version of this dataset is continuously evolving.

| Species | Common Name | Taxonomy |
| --- | --- | --- |
| *Aedes aegypti* | Yellow fever mosquito | Animal |
| *Anopheles gambiae* | African malaria mosquito | Invertebrate |
| *Apis mellifera scutellata* | Honeybee |  |
| *Bombyx mandarina* | Silkworm |  |
| *Caenorhabditis briggsae* | Roundworm |  |
| ***Caenorhabditis elegans*** | Roundworm |  |
| *Culex pipiens* | Common house mosquito |  |
| ***Drosophila melanogaster* R5** | Fruit fly |  |
| *Drosophila melanogaster* R6 | Fruit fly |  |
| *Drosophila pseudoobscura* | Fruit fly |  |
| *Heliconius melpomene melpomene* | Postman butterfly |  |
| *Bos taurus* | Cow | Animal |
| *Canis lupus* | Wolf | Vertebrate |
| *Cynoglossus semilaevis* | Tongue sole |  |
| ***Danio rerio*** | Zebrafish |  |
| *Equus ferus przewalskii* | Prewalskii’s horse |  |
| *Ficedula albicollis* | Collared flycatcher |  |
| *Gallus gallus* | Chicken |  |
| *Gasterosteus aculeatus* | Stickleback |  |
| *Homo sapiens* | Human |  |
| *Lepisosteus oculatus* | Spotted gar |  |
| *Macaca mulatta* | Rhesus macaque |  |
| *Meleagris gallopavo* | Turkey |  |
| ***Mus musculus castaneus*** | House mouse |  |
| *Oryzias latipes* | Medaka |  |
| *Ovis canadensis* | Bighorn sheep |  |
| *Papio anubis* | Olive baboon |  |
| *Sus scrofa* | Wild boar |  |
| *Citrus reticulata* | Mandarin Orange | Plant |
| *Gossypium raimondii* | New world cotton | Woody |
| *Populus trichocarpa* | Black cottonwood |  |
| *Prunus davidiana* | David’s peach |  |
| *Arabidopsis thaliana* | Thale cress | Plant |
| *Brachypodium distachyon* | Purple false brome | Herbaceous |
| *Capsella rubella* | Pink Shepherd’s Purse |  |
| *Citrullus lanatus lanatus* | Watermelon |  |
| *Cucumis sativus* var. *hardwickii* | Cucumber |  |
| *Glycine soja* | Wild soybean |  |
| *Medicago truncatula* | Barrel medic |  |
| *Oryza rufipogon* | Wild rice |  |
| *Setaria italica* | Foxtail millet |  |
| *Sorghum bicolor* subsp. *verticilliflorum* | Wild Sudan grass |  |
| ***Solanum lycopersicum*** | Domesticated tomato |  |
| *Zea mays* ssp *parviglumis* | Teosinte |  |

**Table S5.** Comparing BREC with similar widely used tools. BREC’s provided features and functionalities are compared along with the Recombination Rate Estimator [22] and the MareyMapOnline [21], following a chronological order (the oldest first).

| Features / Tool |  | Recombination Rate Calculator (RRC) | MareyMap Online | BREC |
| --- | --- | --- | --- | --- |
| Publication year |  | 2010 | 2017 | 2020 |
| Genome-specific |  | D. melanogaster | all | all |
| interpolation method | Polynomial | yes | yes | no |
|  | Loess | no | yes | yes |
|  | Cubic spline | no | yes | no |
| Data cleaning |  | no | manually | automatically |
| Data Quality Control |  | no | no | yes |
| Chromatin boundaries identification |  | no | no | yes |
| Software | R package | no | yes | yes |
|  | Web-based GUI | Perl CGI | Shiny | Shiny |

